# High Resolution Single Cell Maps Reveals Distinct Cell Organization and Function Across Different Regions of the Human Intestine

**DOI:** 10.1101/2021.11.25.469203

**Authors:** John W. Hickey, Winston R. Becker, Stephanie A. Nevins, Aaron Horning, Almudena Espin Perez, Roxanne Chiu, Derek C. Chen, Daniel Cotter, Edward D. Esplin, Annika K. Weimer, Chiara Caraccio, Vishal Venkataraaman, Christian M. Schürch, Sarah Black, Maria Brbić, Kaidi Cao, Jure Leskovec, Zhengyan Zhang, Shin Lin, Teri Longacre, Sylvia K. Plevitis, Yiing Lin, Garry P. Nolan, William J. Greenleaf, Michael Snyder

**Author notes:** These authors contributed equally (John W. Hickey, Winston R. Becker, Stephanie A. Nevins).

## Abstract

The colon is a complex organ that promotes digestion, extracts nutrients, participates in immune surveillance, maintains critical symbiotic relationships with microbiota, and affects overall health. To better understand its organization, functions, and its regulation at a single cell level, we performed CODEX multiplexed imaging, as well as single nuclear RNA and open chromatin assays across eight different intestinal sites of four donors. Through systematic analyses we find cell compositions differ dramatically across regions of the intestine, demonstrate the complexity of epithelial subtypes, and find that the same cell types are organized into distinct neighborhoods and communities highlighting distinct immunological niches present in the intestine. We also map gene regulatory differences in these cells suggestive of a regulatory differentiation cascade, and associate intestinal disease heritability with specific cell types. These results describe the complexity of the cell composition, regulation, and organization for this organ, and serve as an important reference map for understanding human biology and disease.

## Introduction

The human adult intestinal system is a complex system of organs consisting of approximately 22 ft of small intestine and 7 ft. of large intestine. This system completes the digestive process begun in the oral cavity and stomach, first absorbing water and small molecule nutrients (e.g. sugars, monovalent ions, and amino acids) in the small intestine, then accumulating larger molecules such as fiber in the large intestine, which serves as an anaerobic fermentation chamber enabling the further breakdown and absorption of these complex molecules and the synthesis, often through alimentary gut microbiota, and absorption of others nutrients such as vitamins^1^.

The small intestine itself is phenotypically heterogeneous, consisting of three morphologically distinct regions: the duodenum, the jejunum and the ileum^2^ . The large intestine can likewise be partitioned into regions including the ascending, transverse, descending, and sigmoid regions. Each of these anatomical regions contains an immense diversity of phenotypically and morphologically distinct cell types. Epithelial, stromal, nerve, and immune cells, each composed of multiple cell types, reside throughout the intestine; immune cells are of particular interest, as they interact with the microbiome and foreign material present in the gut^3^. Although these broad cell types are common to all portions of the intestinal system, prevalences of specific cell types are known to display locational differences: for example Paneth cells are known to populate the small intestine and enteroendocrine L cells are found primarily in the ileum and large intestine ^4, 5^. Moreover, these cell types are spatially organized into different “neighborhoods” across these intestinal regions, with both the composition of these neighborhoods and molecular phenotypes of underlying cellular types varying in relatively unknown ways across these anatomical regions. These differences in both the composition of functional neighborhoods and molecular identity of the cell states that comprise these neighborhoods promise to further reveal the logic by which the human intestine is constructed.

Below we map many portions of the intestine at single cell resolution using single nuclear RNA, open chromatin, and spatial proteomic imaging technologies. Previous studies have mapped cell types using single cell RNA sequencing (scRNA-Seq) and have catalogued cell types across the intestine^6^. We extend this work by the spatial mapping of cells and proteins using CODEX (CO-Detection by indEXing^7–10)^ to characterize approximately 1.2 million cells spatially as well as the mapping of gene regulatory information using single-cell assay of open chromatin using snATAC-seq^11^. We define the relative abundance of distinct cell types across the intestine including the enormous complexity of epithelial cells across different regions of the intestine, and the organization of cells into different multicellular structural niches. We also map gene regulatory differences in these cells suggestive of a regulatory differentiation cascade. These results provide important insights into cell function, regulation, and organization for this complex organ and serve as an important reference for understanding human biology and disease.

## Results

### Mapping the human intestine at single cell resolution

We mapped the cell composition, regulatory information, and spatial distribution of single cells across the intestine of multiple donors using single nuclear RNA-Seq (snRNA-seq), which measures nuclear RNA transcripts in individual nuclei, single nuclear ATAC-Seq (snATAC-seq), which measures open chromatin in single cells, and CODEX, which stains the same tissue section with up to 54 antibody probes against different proteins. We analyzed eight sections from four individuals: three European-ancestry (one male and two female) and one African American male. Age ranges were from 24 to 78 years. The eight regions (in order of trajectory from the stomach) were: duodenum, proximal jejunum, mid-jejunum and the ileum from the small intestine, and ascending, transverse, descending and sigmoid regions of the large intestine. The organs were procured from organ donors and were deemed to be of high quality by RNA quality measures (Methods).

### CODEX Multiplexed Imaging of the Small Intestine and Colon

To create a spatial map of the intestine across the eight different regions, we used CODEX multiplexed imaging. CODEX data retains spatial coordinates of the cell types, thus enabling insight into cellular interactions, composition of multicellular tissue units, and spatial relationships to the overall function of the intestine ^9, 10^. We first validated and optimized CODEX staining, imaging, and image processing for fresh frozen samples on one subject (B001). Four different regions of the colon and small intestine were combined on the same coverslip to minimize batch effects (Fig. 1A). Using a 44 marker antibody panel we stained and detected immune, stromal, and epithelial cells and multicellular neighborhoods from approximately 16 mm^2^ sections (Supplemental Fig. 1-3). Three epithelial, six immune, and six stromal cell types were identified— resulting in two epithelial, three immune, and five stromal multicellular neighborhoods characteristic of the intestine (e.g., immune follicle).

**Figure 1:**
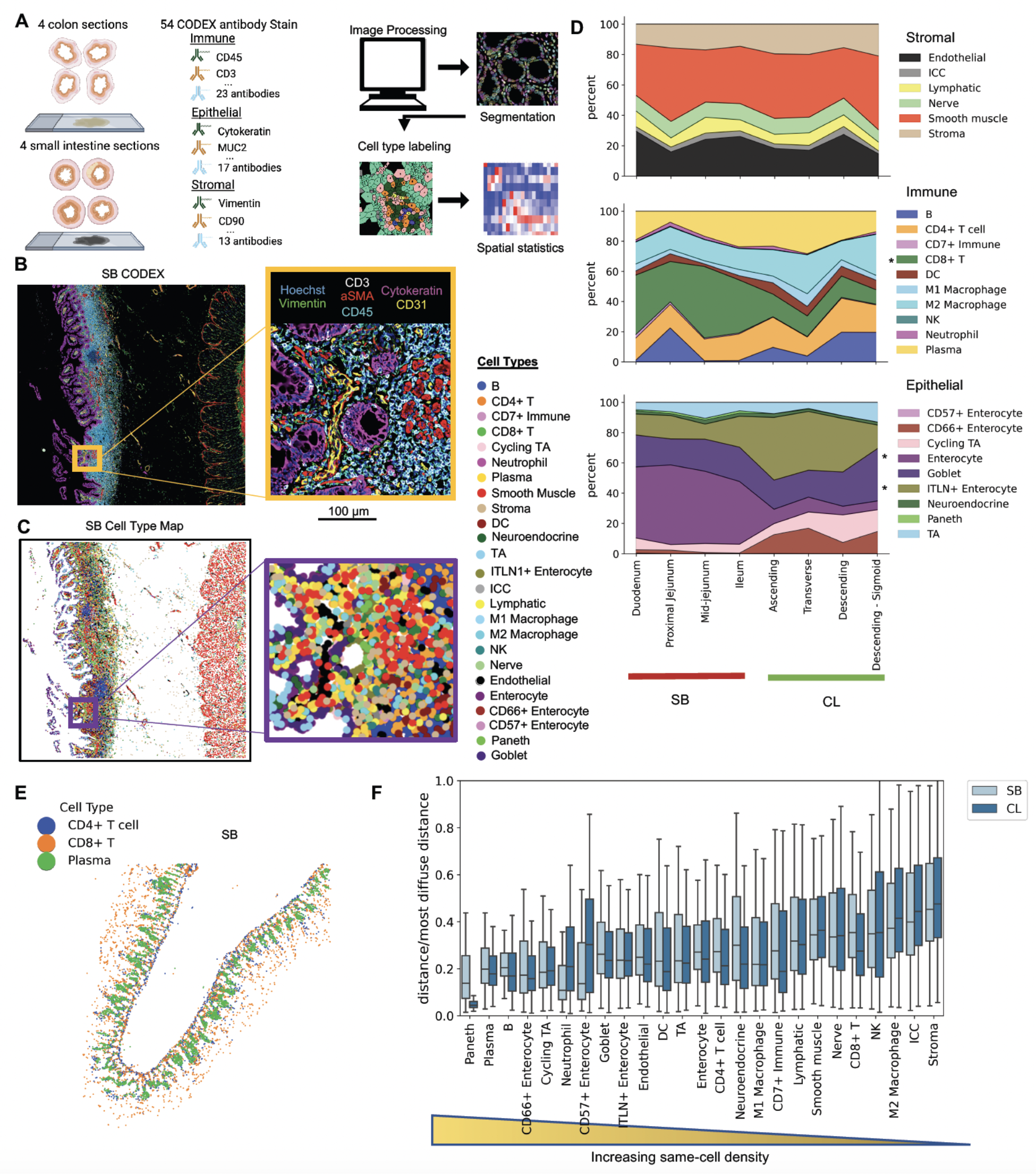
CODEX multiplexed imaging of 8 regions from the small intestine and colon to create a single-cell map of the healthy human intestine. A) Schematic for how CODEX multiplexed imaging was performed on arrays of 4 different sections of either colon and small intestine from the same donor simultaneously. Image processing steps done to extract single-cell spatial data. B) An example CODEX fluorescent image of one region of the small bowel (SB) for CODEX with 6/54 markers shown for one donor with C) accompanying cell type map following cell segmentation and unsupervised clustering. D) Cell type percentages from CODEX data averaged across 3 donors stained with an updated CODEX antibody panel. Cell types are normalized by stromal, immune, and epithelial compartments. Asterix indicates p-value less than 0.05 difference in cell type percentage from the SB to the colon (CL). E) One representative cell type map with only plasma cells, CD4+ T cells, and CD8+ T cells shown. F) Quantification of the same-cell density that is measured as an average distance of its 5 nearest same-cell neighbors divided by the most diffuse same-cell distance.

**Figure 2:**
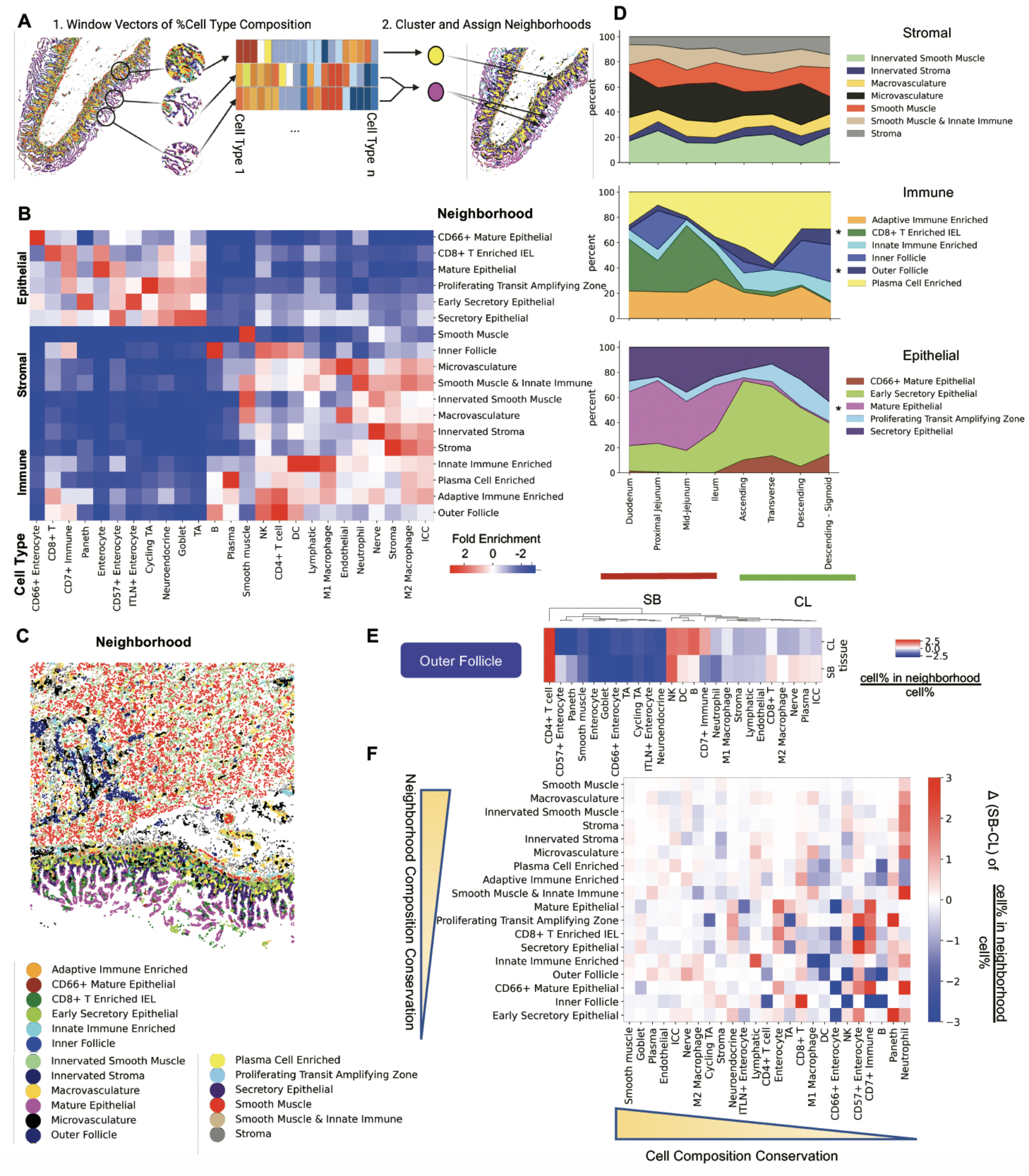
Multicellular neighborhood analysis of intestine. A) Neighborhood analysis was done by taking a window across cell type maps and vectorizing the percentage of cell types in each window, clustering windows, and assigning clusters as cellular neighborhoods of the intestine. B) 18 unique intestinal multicellular neighborhoods (y axis of heatmap) were defined by enriched cell types (x axis of heatmap) as compared to overall percentage of cell types in the samples with C) an example of where neighborhoods are mapped back to the tissue to show overall tissue structures. D) Neighborhood percentages from CODEX data averaged across 3 donors normalized by stromal, immune, and epithelial compartments. Asterix indicates p-value less than 0.05 difference in cell type percentage from the small bowel (SB) to the colon (CL). E) Heatmap of enriched cell types for just the *outer follicle* neighborhood compared between SB and CL. F) Difference in composition in neighborhood by cell type for all neighborhoods based on subtracting the fold enrichment in SB from CL.

**Figure 3:**
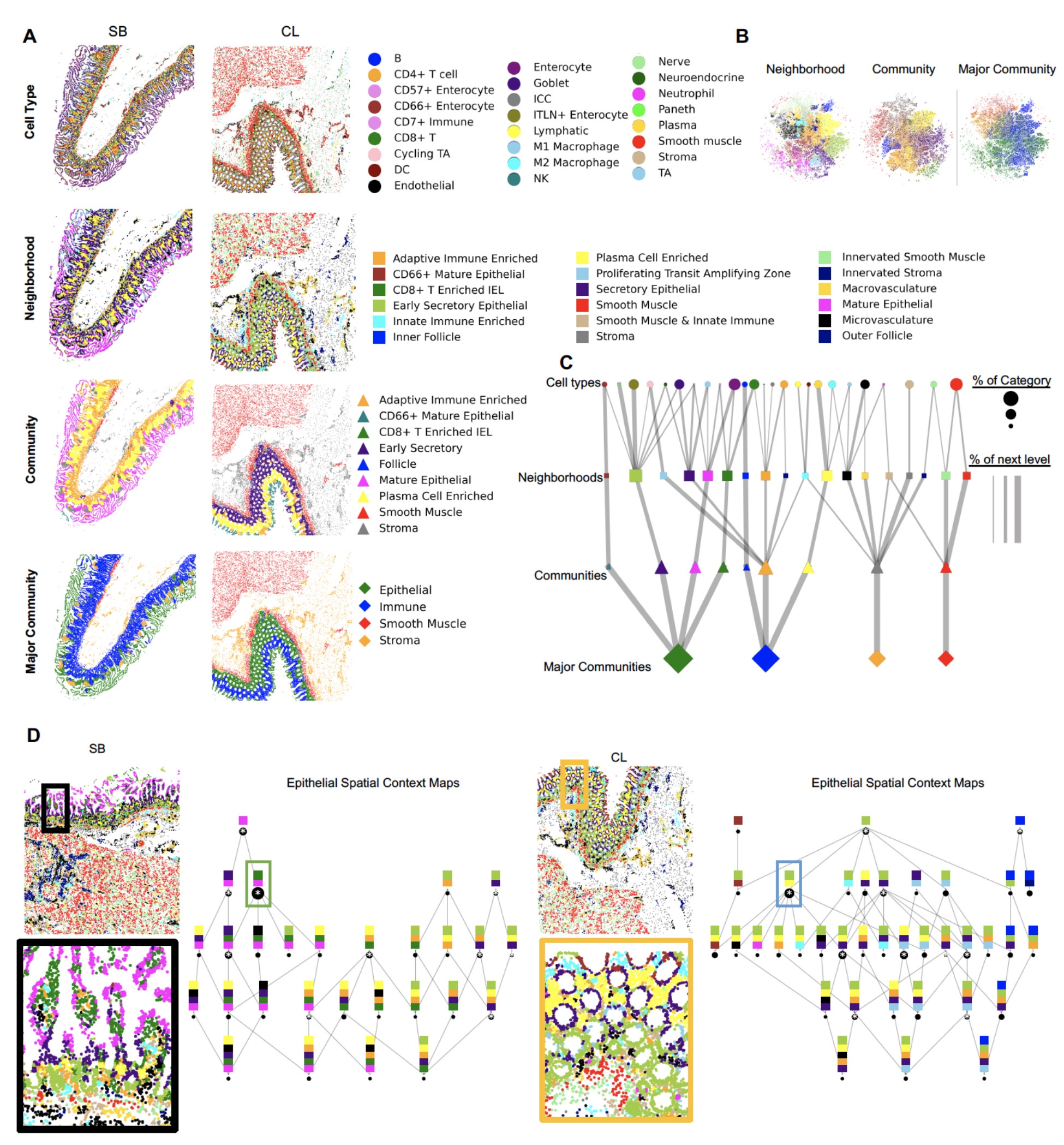
Multi-level hierarchical structural description of small intestine and colon. A) Representation of multiple levels of hierarchical description: i) cell type, ii) multicellular neighborhood, iii) community (based on clustering windows of cell neighborhoods), and iv) major community (major types of communities) compared for the small bowel (SB) to the colon (CL). B) tSNE of neighborhood vectors with neighborhood, community, and major community labels projected. C) Graph of multi-level structure of the tissue as broken down by the different structures. Shapes correspond to structural level, colors represent individual categories, size of shapes represents the percent contribution to tissue, and the size of connected lines represents the overall contribution to the next level of structure as moving down the graph in increasing tissue structural hierarchy. D) Spatial context maps of SB and CL highlighting relationships of multicellular neighborhoods. This structure is defined by the number of unique neighborhoods required to make up at least 85% in a given window. This graph was constructed from only the epithelial and immune regions as defined by the major community labels of cells. Circles represent the number of cells represented by a given structure.

Epithelial cell subtypes are known to be spatially restricted and key to overall function of the gut^12^; therefore we expanded our CODEX antibody panel by adding and validating 17 intestine-specific markers (Supplemental Information 1). We similarly imaged the eight region samples from the three additional donors with this updated 54-antibody panel (Fig. 1B, Supplemental Fig. 4) which enabled the identification of an additional 10 cell types, such as Paneth and goblet cells (Fig. 1C, Supplemental Fig. 5,6). We propagated these cell type labels to two additional donors using the geometric deep learning method STELLAR that we developed for multiplexed spatial data (*Brbic et. al,* co-submitted).

**Figure 4:**
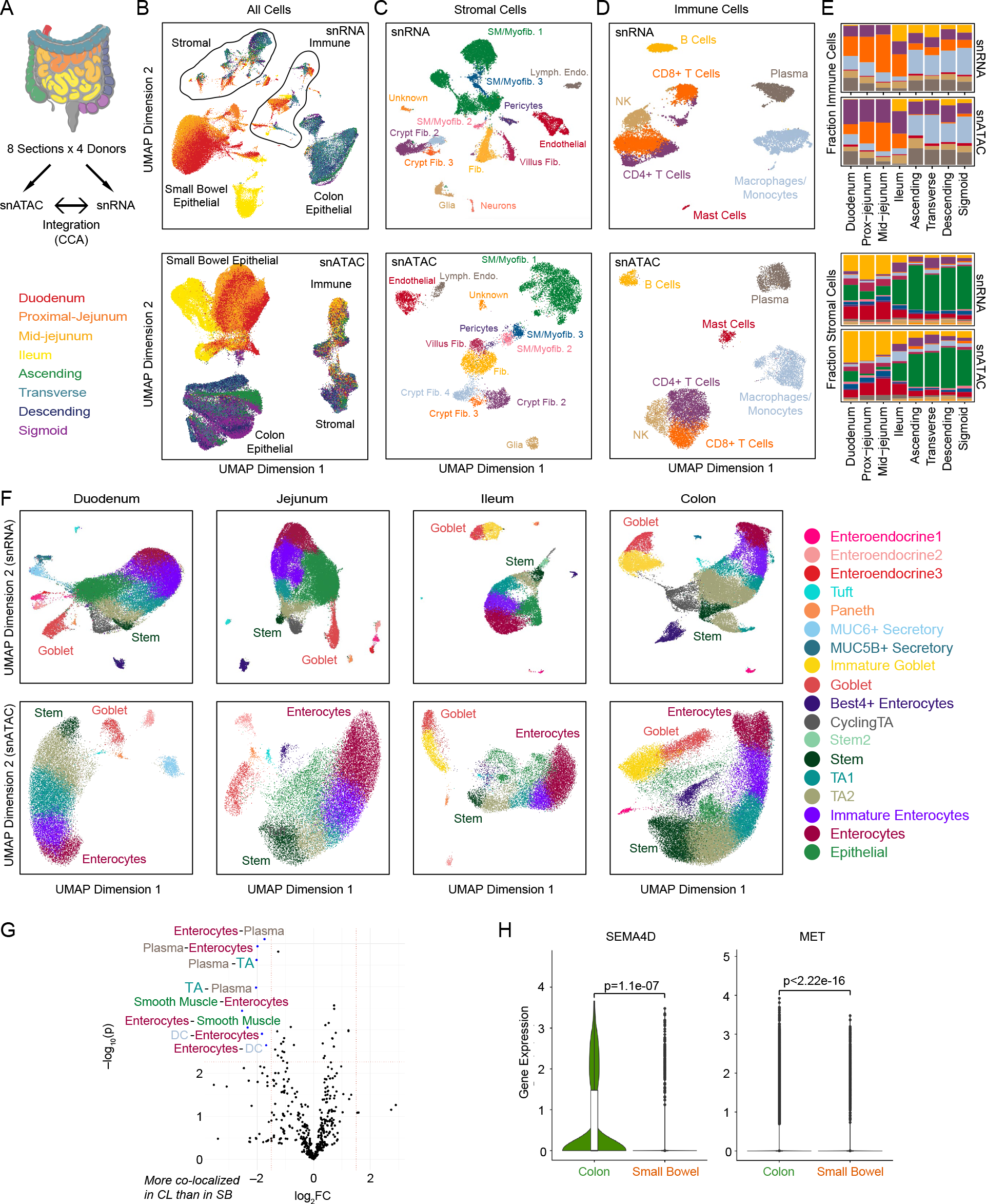
Single-cell atlas of gene expression and chromatin accessibility in the human intestine. A) Sections of the intestine from which cells were isolated for snRNA-seq and snATAC- seq. B) UMAP representation of all cells colored by the region of the intestine that they were isolated from as indicated in A. C, D) UMAP representation of all stromal (C) and immune (D) cells colored by cell type. E) Stacked bar plot representation of the fraction of total immune (top) and stromal (bottom) cells isolated from each region consisting of each cell type. F) UMAP representation of epithelial cells in the 4 primary regions of the intestine. Jejunum includes both proximal- and mid-jejunum samples. Colon includes samples from ascending, transverse, descending, and sigmoid colon. For all UMAP representations, snRNA is on the top and snATAC is on the bottom. G) Large colon (CL) and small bowel (SB) show differences in cell-cell co- localization patterns; annotated cell-pairs are more colocalized in the colon compared to the small bowel. H) SEMA4D ligand expression in plasma cells and MET receptor gene expression in TA2 cells, showing higher expression in colon than small bowel.

**Figure 5:**
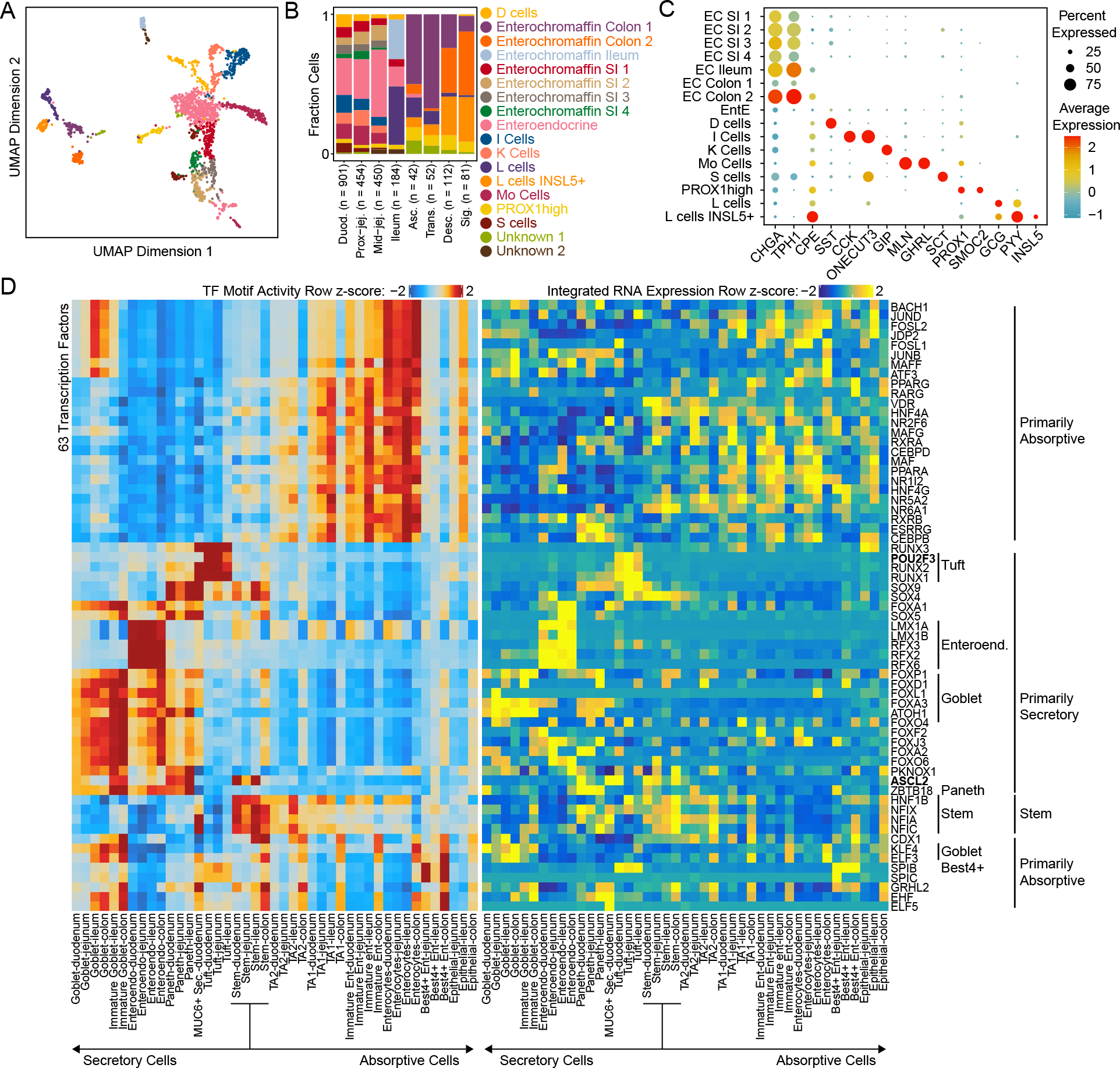
Regulatory transcription factors in the human intestine. A) Sub-clustering of enteroendocrine cells from all regions of the intestine. Cells are colored by cell types as defined in B. B) Fraction of all enteroendocrine cells in each region of the small intestine and colon made up of each enteroendocrine/enterochromaffin subtype. C) Dotplot representation of the expression of subtype specific enteroendocrine and enterochromaffin marker genes in different enteroendocrine cell types in our datasets. D) Heatmap representation of transcription factors whose integrated gene expression was correlated with their motif activity in one region of the intestine. Row z-scores of ChromVar deviation scores are shown on the left and row z-scores of integrated TF expression are shown on the left.

**Figure 6:**
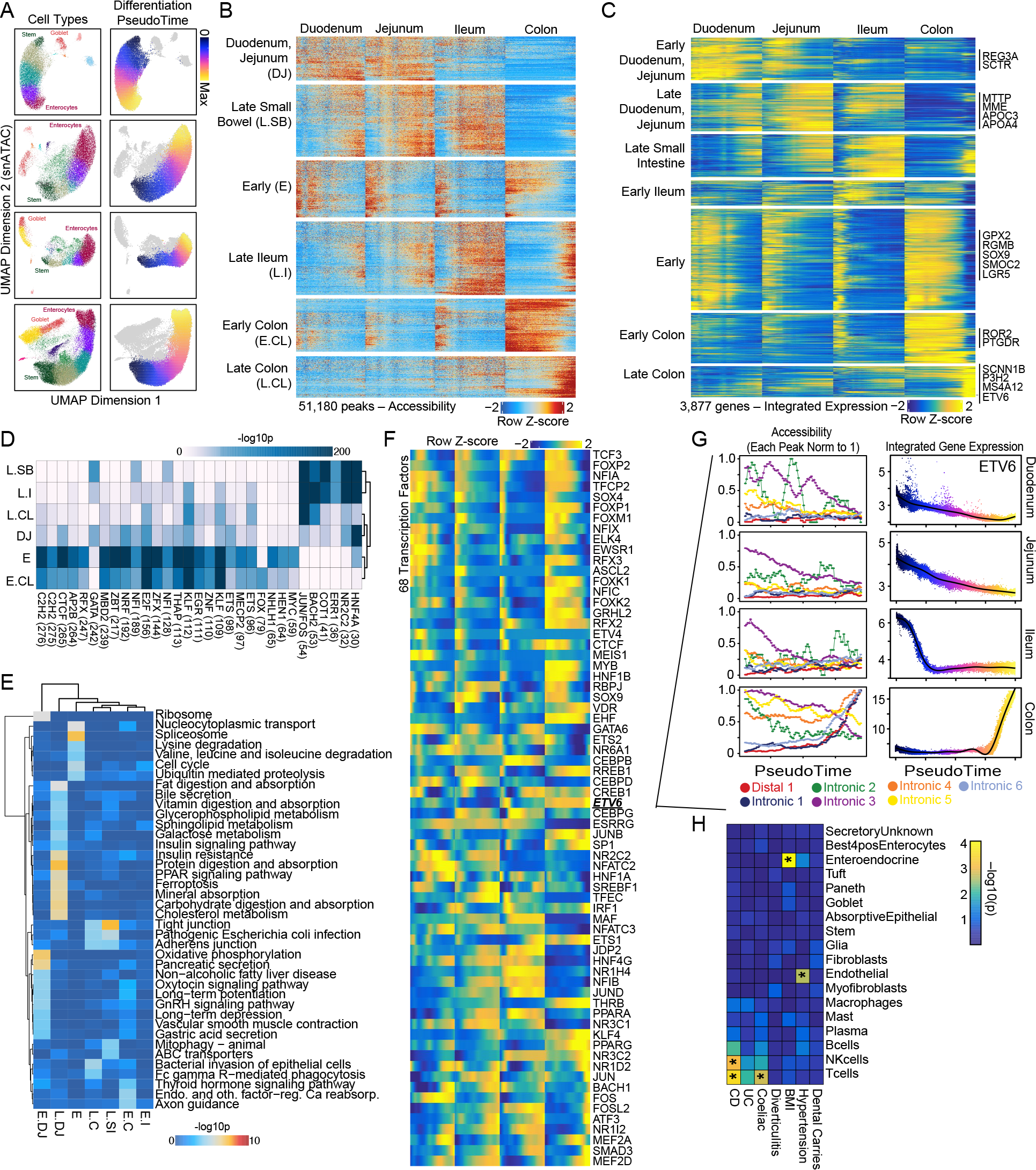
Regulation of differentiation in the human intestine. A) UMAPs depicting the cells in the four primary regions of the intestine (duodenum, jejunum, ileum, and colon), labeled by cell type (left) and differentiation pseudotime (right). B, C) Variable peaks (B) and genes (C) identified along the absorptive differentiation trajectories. The rows represent the row z-scores of accessibility for each peak or expression for each gene. Columns represent the position in the pseudotime from start to end for each section of the intestine. Peaks and genes are k-means clustered and the clusters are labeled based on the dominant time and location where they are most accessible/expressed. D) Hypergeometric enrichment of motifs in the clusters of peaks identified in B. E) Enrichment of KEGG pathways in the clusters of genes identified in C. F) Integrated gene expression of TFs whose gene expression is correlated with ChromVar motif activity along the differentiation trajectory. G) Accessibility at peaks correlated with the expression of ETV6 along the differentiation trajectory in each region is plotted on the left. Each peak is normalized to the maximum accessibility along any of the trajectories. Integrated gene expression of ETV6 along the differentiation trajectory in each region is plotted on the right. H) LD Score Regression to identify enrichment of GWAS SNPs in cell-type specific marker peaks. Unadjusted p-values are plotted in the heatmap. Significance is indicated by an asterisk in each box, as determined by a Bonferroni-corrected p-value of less than 0.05. P-values for determining significance were adjusted for the number of cell classes tested.

We used this resultant dataset to compare the cellular composition and organization across the different tissue regions. Cell types were separated by immune, stromal, and epithelial groupings because tissue sections had unequal representation of epithelial or stromal components (Fig. 1D, Supplemental Fig. 8). We observed no statistical differences in the representation of cells in the stromal compartment, where we would not expect to see major differences in vascularization, innervation, or muscle layers within our resolution of cell types. In the immune compartment, we observed a decrease in CD8+ T cells from the small intestine to the colon, consistent with previous flow cytometry-based observations^13^. Similarly, we observed a decrease in absorptive enterocytes, an increase in secretory enterocytes (goblet and immature goblet cells), and an absence of Paneth cells when moving from the small intestine to colon. Cell type compositions across the different regions of the small intestine and colon differ more than across donors (for the same region), providing confidence in the reproducibility of multiplexed imaging results (Supplemental Fig. 9).

Cellular density reflects whether a cell has broad functions over large regions, is spatially restricted for specialized functions, or has need for specific cell-cell interactions. Moreover, aberrations in cellular density have been associated with disease states such as inflammatory bowel disease (IBD). As the intestine is the largest immune organ in the body^14^, evaluating immune cell localization and cellular interactions in the gut is critical for understanding oral vaccine design^15^, interactions regulating the gut microbiota^16^, regulation of allergic food responses^17^, axes critical for wound repair^18^, and immune system responses and immunotherapies for cancer^19–22^.

Examination of three immune cell types (CD4+ T, CD8+ T, and plasma cells) revealed gross differences in cellular density (Fig. 1E). Visual inspection suggested that plasma cells had the highest same-cell type density, followed by CD4+ T cells, then CD8+ T cells, which were most diffuse through mucosa areas. We quantified this observation by calculating the average distance of a given cell type to its five nearest neighbors of the same cell type and dividing by the distance expected if all the same cells were arranged homogeneously across the tissue (Fig. 1F). Consequently, scores closer to 1 represent lower same cell density. Quantification confirmed our qualitative observations, showing that plasma cells were most dense (∼0.2), followed by CD4+ T cells (∼0.3), then CD8+ T cells (∼0.37). Support functions from CD4+ T cells are largely restricted to immune cell-cell interactions, whereas for immune surveillance of CD8+ T cells we would expect more diffuse distribution. Interestingly, these densities were largely conserved across the intestine between similar cell types. These results suggest an important role for spatial restriction of immune cell subtypes along the length of the intestine.

### Multicellular Neighborhood Analysis of the Intestine

To provide a global view of intercellular interactions, cellular densities, and overall multicellular structures of the intestine, we performed cellular neighborhood analysis^10^. Briefly, this analysis involved (i) taking windows of cells across the entire cell type map of a tissue with each cell as the center of a window, (ii) calculating the percentage of each cell type within this window, (iii) clustering these vectors, and (iv) assigning overall structure based on the average composition of the cluster (Fig. 2A). Cellular neighborhood analyses on the CODEX data of the intestine revealed 18 significant multicellular structures with major epithelial, stromal, and immune based neighborhoods (Fig. 2B, C; see methods).

Congruent with our observation of high same-cell density of plasma cells (Figure 1), we observed a *plasma cell enriched* neighborhood driven by increased density of plasma cells (Fig. 2B). This *plasma cell enriched* neighborhood also exhibits co-enrichment of CD4+ T cells and antigen-presenting cells such as dendritic cells and macrophages. These observations are consistent with recent work suggesting that CD28 engagement on plasma cells from antigen-presenting cells within the bone marrow can maintain long-term survival in plasma-specific niches^23, 24^. Furthermore, antigen-presenting cells are sources of APRIL, which drives ectopic germinal center formation and plasma cell infiltration in cases of inflammation^25^. Collectively, these observations suggest a role for antigen-presenting cells in curating a subepithelial niche for plasma cells in the intestine.

Interestingly, despite low same-cell density relative to other cells in the intestine, CD8+ T cells were conserved and enriched in two major neighborhoods (Fig. 2B). One of these neighborhoods (*CD8+ T enriched IEL*) exhibits enrichment of both epithelial cell types and CD8+ T cells. These CD8+ T cells are commonly referred to as intraepithelial lymphocytes (IELs), which are “soldiers on the front lines,” critical for rapid immunological responses, protection against infection (through MHCI binding and cytotoxicity^26^), and maintenance of epithelial integrity (via regulation of epithelial cell proliferation^27^). Dysregulation of these cells is associated with IBD and celiac disease. However, due to challenges in isolating these T cells, they have been difficult to study. This *CD8+ T enriched IEL* neighborhood was one of the few neighborhoods to change in prevalence from the small intestine (∼30%) to the colon (∼3%) (Fig. 2C, Supplemental Fig. 9). This variation may reflect differences in antigen exposure, as cells in the small intestine have increased exposure to dietary antigens, whereas cells in the colon likely experience more exposure to gut microbiota.

CD4+ T cells contributed to 5 diverse multicellular neighborhoods of the intestine, including *inner follicle, innate immune enriched, plasma cell enriched, adaptive immune enriched,* and *outer follicle* (Fig. 2B). This broad neighborhood membership is fitting, given that CD4+ T cells coordinate innate and adaptive immune responses. CD4+ T cells, B cells, and DCs (dendritic cells) membership defined two different follicle-based structures. The first of these structures, which exists in outer regions of the follicle, exhibited higher enrichment of CD4+ T cells, while inner regions of the follicle were enriched in B cells (Fig. 2B). The presence of the *inner follicle* (i.e., the germinal center) neighborhood was dependent on a fully mature lymphoid follicle like a Peyer’s patch within the image (Fig. 2D). However, the *outer follicle* neighborhood was constant across the intestine and defined by B cells, DCs, and CD4+ T cells that were enriched in *outer follicle* neighborhoods compared to *inner follicle* structures. These data agree with previous work defining a continuum of lymphoid tissues within the intestine, where smaller structures like cryptopatches have the potential to develop into larger follicles^28–30^ especially in cases of inflammation^31^.

Previous research has also indicated differences in lymphoid follicle development in the small intestine vs the colon^32, 33^. By comparing the *outer follicle* neighborhood cell type compositions in the small intestine versus the colon (Fig. 2E, Supplemental Fig. 10A, B), we observed that while both are driven by an enrichment of CD4+ T cells, the colon had a higher enrichment of B cells and DCs, whereas the small intestine was enriched for CD8+ T cells, nerves, and plasma cells. This structural similarity is identified by the neighborhood analysis, but major differences of cell compositions within these similar neighborhoods can also be identified.

We next compared the relative fold enrichment of all neighborhood compositions across the small intestine and colon, to create a heatmap of neighborhood differences (Fig. 2F). This heatmap is ranked both in terms of greatest total difference within both neighborhood and cell type categories, indicating neighborhoods with greatest conservation across the intestine, as well as the cell types that are most conserved across neighborhoods. Multicellular structural conservation across the intestine suggests a required function by a set of cell types, whereas less conservation indicates a core functionality as well as a need for compositional flexibility based on anatomical location. We found that both inner and outer follicle structures are less conserved, whereas stromal neighborhoods are more conserved. We also observe that the *early secretory epithelial* neighborhood, which contains Paneth cells in the small intestine and not in the colon, is the least conserved neighborhood—indicating different crypt microenvironments across the intestine (Fig. 2F, Supplemental Fig. 10C).

### Hierarchical Structural Analysis of Intestine and Mucosal Epithelial Compartment

Multicellular neighborhood analysis revealed key differences in structural composition across the intestine as well as composition of these neighborhoods. However, this analysis did not reveal how these multicellular neighborhoods interact with one another, or how they are spatially structured in the tissue. Understanding how multicellular groups are related is key to both defining the hierarchy of tissue organization, as well as defining key functional tissue interfaces.

We investigated higher order structural organization using several methods. First, we clustered windows of neighborhood compositions in a manner similar to our methods for defining the neighborhoods from cell types; this generated *communities* and *major communities* of neighborhoods (Fig. 3A, Supplemental Fig. 11), and revealed spatial layering of the intestine, moving from the smooth muscle, stroma, and within epithelial areas. Interestingly, comparing the localization of *immune major communities* between the small bowel and colon showed differential localization localizing to the base of the epithelium compared to the large intestine located towards the apex, likely reflective of differential microbial loads. Moreover, tSNE representations of the cell neighborhood vectors recapitulated overall tissue structure; for example, major community maps show smooth muscle and stromal neighborhoods in close proximity, abutted by immune neighborhoods located between epithelial structures (Fig. 3B).

To relate the various levels of spatial organization to one another we created a hierarchical structure network graph (Fig. 3C, Supplemental Fig. 12). Each level of this graph is connected to the next by its major contributors to these higher-order structures. Using this intuitive formalism, we observe crosstalk between stromal and smooth muscle cell types and structures, which are in turn isolated from epithelial and immune components that are more entwined with one another. Additionally, major communities of epithelial and immune origin are much more diverse in composition across the levels. Using this graph structure, we can also observe multi-level relationships between the structures. For example, the *adaptive immune enriched* community is an important intersection of multiple neighborhoods: *proliferating transit amplifying zone, secretory epithelial, adaptive immune enriched, outer follicle*, and *innate immune enriched*. Consequently, this indicates that these important immune enriched neighborhoods, that have yet to become larger structured immune follicles, are poised with specialized niches flanking the secretory and transit amplifying zones of the intestine.

We leveraged this hierarchical deconstruction of tissue organization to examine interactions in multicellular neighborhoods^34^. We isolated the cells as defined by a major community of *epithelial* and *immune* organizations (Fig 3A), and used a similar principle to our neighborhood analysis (Fig. 2A) to extract interaction information. We quantified the composition of windows across the neighborhood maps based on the percentage of neighborhoods in each window. Then instead of clustering window composition vectors, we evaluated which combination of neighborhoods made up at least 85% of the neighborhoods in each window, summed the unique neighborhood combinations, and connected these structural interactions in a network graph structure for both small intestine and colon (Fig. 3D).

From this analysis, already noted differences in composition are clear, where the *mature epithelial* and *CD8+ T enriched IEL* neighborhood combination is present in the small intestine but nonexistent in the colon (Fig. 3D, green box). In the colon the *early secretory epithelial* neighborhood is enriched combined with several other neighborhoods including the *proliferating transit amplifying zone* and *plasma cell enriched* (Fig. 3D blue box). In addition, the *plasma cell enriched* neighborhood is connected to many neighborhoods, but it is often co-enriched with the *adaptive immune enriched* neighborhood, which is also shared by both small intestine and colon graphs. The interconnectedness of the *plasma cell enriched* neighborhood can also be seen from the order of neighborhoods in the mucosal regions in high resolution images (Fig. 3D, black/orange boxes). Early secretory and adaptive immune neighborhoods are enriched at the bottom of the crypt followed by secretory and plasma cells enriched through mid-parts of the crypt followed by innate immune and mature epithelial neighborhoods. In conclusion, our hierarchical mapping data further confirms compositional differences in multicellular structures between small intestine and colon but also highlights conserved multicellular structure interactions and an important distribution of distinct cell types in subregions of the intestine.

### Single-cell transcriptomic and chromatin analyses reveal cell composition differences across regions of the intestine

CODEX revealed distinct compositions of cells and their arrangement across different intestinal regions. However, the CODEX experiments included only 54 probes, potentially limiting both the number and complexity of cell types that can be identified and their distribution in different regions of the intestine. To overcome these limitations as well as create a gene-regulatory atlas of the human intestine, we generated 156,119 snRNA-seq and 153,997 snATAC-seq profiles on the eight intestinal regions from the four donors (Figure 4A, Methods, Supplemental Fig. 13A,B).

To examine global differences between regions of the intestine, we first clustered cells from all regions of the intestine and found that immune and stromal cells from all regions clustered together whereas epithelial cells separated based on whetherg they were isolated from 1) duodenum or jejunum, 2) ileum, or 3) colon (Figure 4B). We next subclustered immune cells, stromal cells, and epithelial cells from different regions of the intestine, revealing a total of 7 immune, 13 stromal, and 18 epithelial cell-types (Figures 4C, 4D, and 4F). Cell-types were annotated by examining gene expression levels and gene activity scores of known marker genes as well as by labeling the datasets with previously published scRNA-seq data^35^ (Methods).

Within the immune compartment, we identified clusters of CD4+ and CD8+ T-cells (CD2, CD3E, IL7R, CD4, CD8), B-cells (PAX5, MS4A1, CD19), Plasma cells (IGLL5, AMPD1), NK cells (SH2D1B), macrophages/monocytes (CD14), and mast cells (HDC, GATA2, TPSAB1) in both the snRNA and snATAC datasets (Figures 4C and Supplemental Fig. 13C). NK and T-cells from one donor clustered separately from the other T-cells in the snRNA data, likely because this donor was much younger (24 years) than other donors in this study (Supplemental Table S1). The majority of the immune cell types were identified in both the single-nuclei data and CODEX data, with the exception of Mast Cells which were only identified in the single-nuclei data, and dendritic cells and neutrophils which were only found in the CODEX data. Similar to the results of the CODEX experiments (Figure 2E), some immune cell types were differentially abundant along the intestine. For example, T cells were more prevalent in the small intestine, whereas B cells and macrophages/monocytes were more frequent in the colon (Figure 4E).

Within the stromal compartment, we annotated eight fibroblast subtypes, glial cells, neurons, and endothelial cells (Figure 4D). Cells with high expression of MYH11 and ACTA2 were classified as smooth muscle/myofibroblasts. We also identified fibroblasts with high levels of WNT agonists, such as RSPO3, which are thought to be present at the crypts, and fibroblasts with high expression of WNT5B and BMP transcripts thought to be present at the villi^35, 36^ (Supplemental Fig. 13D). Similar to the immune cells, we observed changes in cell type abundance along the intestine (Figure 4E). For example, the smooth muscle/myofibroblasts were least abundant in the duodenum and jejunum, had intermediate abundance in the Ileum, and were most abundant in the colon. Conversely, villus fibroblasts and endothelial cells exhibited the opposite trend; they were most abundant in the duodenum and jejunum, less abundant in the Ileum, and least abundant in the four regions of the colon (Figure 4E).

Given that epithelial cells initially clustered based on location (Figure 4B), we subclustered and annotated epithelial cells from different primary locations—duodenum, jejunum, ileum, and colon—separately. In annotating the cells, we observed similar cell types within the different intestinal regions. For example a differentiation trajectory from stem cells to mature absorptive cells (enterocytes) was evident in all regions. We divided this differentiation trajectory into five cell types previously defined in other studies (Stem>TA2>TA1>Immature Enterocyte>Enterocyte)^35^. However, we observe that these cells exist along a continuum and therefore the exact number of cell types and locations of the divisions between cell types is arbitrary to some extent, and changing the resolution during clustering results in more or fewer clusters along this trajectory. In addition to the absorptive cells, we also observed Goblet Cells, Best4+ Enterocytes, Tuft Cells, and Enteroendocrine Cells in all regions of the intestine, although there was greater diversity in the Enteroendocrine Cells observed in the small intestine, where multiple discrete clusters (Figure 4E; Enteroendocrine 1–3) were observed. As expected and consistent with the CODEX results, Paneth cells were observed in all regions of the small intestine, but were not found in the colon. Finally, the duodenum contained two additional clusters that were identified in the snRNA and one that was identified in the snATAC datasets. These cells clustered separately from the absorptive cell types, and we found that the kegg pathway Mucin type O−glycan biosynthesis was enriched in the marker genes of one of these cell types (Supplemental Fig. 14A, B). When examining specific marker genes in these clusters, we found that one cluster had high expression of MUC5B and the other cluster had high expression of MUC6 and TFF2 (Supplemental Fig. 14C), suggesting these are likely different types of mucin producing cells, with the MUC6+ cells likely representing the cells of the Brunner’s glands ^37^.

To further explore the diversity of enteroendocrine cells along the human intestine, we subclustered enteroendocrine cells from all regions of the intestine (Figure 5A), and annotated the clusters based on expression of enteroendocrine marker genes (Figure 5C). We identified many known subtypes of enteroendocrine cells including D cells (SST high), I cells (CCK high), K cells (GIP high), Mo cells (MLN high), S cells (SCT high), and L cells (PYY high). However, we also identified multiple clusters of enterochromaffin cells, which express TPH1, an enzyme involved in serotonin synthesis. There was a cluster of enteroendocrine cells that did not express any of these specific markers, which we labeled as enteroendocrine in the subclustered dataset. L cells formed two distinct clusters, one of which has high expression of INSL5, which we labeled as INSL5+ L cells. The absolute number of enteroendocrine and enterochromaffin cells was highest in the duodenum followed by the jejunum (Figure 5B). We observe large shifts in the fraction of each subtype of enteroendocrine cells along the intestine (Figure 5B). For example, D cells, which express SST, were most abundant in the duodenum, as expected from previous studies^38^. In the colon, enterochromaffin and L cells were the most abundant, with some shifts observed along the length of colon, including an increase in INSL5+ L cells in the distal colon. These results provide exceptional detail on the complexity of enteroendocrine and enterochromaffin cells, potentially defining new subtypes of enterochromaffin cells, and describe how their populations change along the length of the intestine.

### Nominating Specific Molecular Interactions in Neighboring Cells

The CODEX data enables the assignment of neighboring cell types and the snRNA-Seq data provides a description of the molecules expressed at the RNA levels in these cells. By combining both data types we can nominate potential ligand receptor pairs that may facilitate cell-to-cell communication for these cell types.

We used the colocalization quotient to measure the degree to which cell type A is preferentially associated with cell type B. We identified significant pairwise cell-type colocalizations and focused on those that were significantly different between small bowel and colon. This analysis identified eight cell type pairs that were more colocalized in colon than small bowel (Figure 4G); six pairs involved enterocytes, plasma cells, TAs and smooth muscle (Supplemental Table 2). Cell type pairs involving plasma cells are more colocalized in the descending-sigmoid colon tissue section than in other sections (Supplementary Fig. 15).

Using snRNA-Seq data from these six cell types we performed differential expression analyses of ligands and receptors (see methods) and identified 48 pairs of ligands and receptors that are more expressed in colon than small bowel (Supplemental Table 3). As an example, we found the ligand SEMA4D and receptor MET upregulated in plasma cells and TA2, respectively, in colon tissue (Figure 4H); this was not observed in the small intestine and was only expressed in 4.1% across all pairs of cell types (Supplemental Table 3). SEMA4D signaling has been associated with a diversity of immunological disorders and plays an important role in cell-cell communication^39^, particularly in B cell aggregation and long-term survival^40^. Implication of the MET receptor on TA2 cells in the colon, compared to TA2 cells in the small intestine, is further evidenced by (i) enrichment of downstream responses to MET activation including RAS and MAPK signalling (Supplemental Fig. 16, Supplemental Table 4), and (ii) upregulation of plexins (Supplemental Fig. 16) known to interact with high affinity to semaphorins and transactivate MET. This possible interaction is consistent with the previous characterizations of the CODEX multiplexed imaging data of distinct plasma-cell enriched multicellular neighborhoods, conservation of plasma-cell enriched neighborhoods across the intestine, and differential connections of these neighborhoods comparing the small intestine to the colon (Figure 3D). Consequently, this indicates a potential differential survival signal that maintains the *plasma-cell enriched* neighborhood in the colon. Overall, these results nominate potential ligand-receptor interactions that mediate specific cell type interactions in distinct regions of the intestine and provide a template for other atlas efforts integrating spatial multiplexed imaging data with paired snRNAseq assays.

### Integration of snRNA and snATAC data nominates transcription factors regulating gene expression in different cell types

To obtain insights into the factors and events that control intestinal differentiation, we next investigated potential transcription factors (TFs) regulating gene expression in different intestinal cell types. We first computed ChromVar deviation scores^41^ for each cell in our dataset, which allows us to identify TF motifs that are associated with chromatin accessibility in different cell types. Because many TFs share similar motifs, examining TF expression in conjunction with motif activity helps identify the specific TFs that are functional in different intestinal cell types. To examine TF expression, we integrated the snRNA and snATAC data using canonical correlation analysis to align the datasets and assign snRNA data to each snATAC cell (Methods)^42, 43^. We next identified the TFs with the highest correlation between their gene expression and the chromatin accessibility activity level of their putative DNA binding motifs^44^ to nominate TFs directly driving accessibility changes. Across the intestine, this analysis revealed 63 TFs with motif activity that strongly correlated with expression (r>0.6; Figure 5D). Broadly, we observe TFs that are active in the secretory lineage and TFs that are active in the absorptive lineage with a relatively small subset of TFs that participate in both (e.g. KLF4 and ELF3 in colon enterocytes and goblet cells throughout the intestine; Figure 5D).

This analysis highlights many TFs known to serve important roles in the intestine. For example, ASCL2, a master regulator of intestinal stem cells^45^, exhibited high expression and accessibility of its motif in stem-cells. Other TFs with high expression and motif activity in Stem Cells include NFIX, NFIA, NFIC, and HNF1B. Within the secretory lineage, POU2F3, a regulator necessary for the development of tuft cells in mice^46^, was highly expressed and had high motif activity in tuft cells throughout the human intestine. Along with POU2F3, RUNX1 and RUNX2 also exhibited high expression and accessibility in tuft cells throughout the intestine. Among goblet cells, KLF4, which is required for terminal differentiation into colonic goblet cells in mice^47^, exhibited high gene expression and motif activity. Surprisingly, expression and motif activity of KLF4 was also high in differentiated absorptive epithelial cells (immature enterocytes and enterocytes) in the colon, but not in other regions of the intestine indicating location-specific regulation. Within enteroendocrine/enterochromaffin cells, LMX1A, LMX1B, RFX2, RFX3, and RFX6, exhibited high expression and accessibility. Of these, LMX1A, has been proposed as a regulator of the enterochromaffin lineage and is a regulator of TPH1, an enzyme involved in serotonin synthesis^48^. When we examine expression of LMX1A in enteroendocrine and enterochromaffin subtypes, we observe that it is expressed only in enterochromaffin cells (Supplemental Fig. S14D). Similarly, RFX6 is a proposed regulator of enteroendocrine cell differentiation, and loss of RFX6 impairs enteroendocrine cell differentiation in mice^49^. Together, these results support previous findings and nominate additional TFs that may be important regulators of distinct intestinal cell types which can vary across the different regions of the intestine (e.g. KLF4).

### Differentiation trajectories across the intestine reveal distinct gene regulation programs

Intestinal stem-cells continuously differentiate into mature enterocytes, goblet cells, and other specialized cell types such as enteroendocrine, tuft, and paneth cells, renewing the epithelial lining approximately every three to seven days^50^. To map the regulatory and gene expression changes that accompany stem-cell differentiation into mature enterocytes, we defined differentiation trajectories along this pathway in the single nuclei data from duodenum, jejunum, ileum, and colon (Figure 6A, Methods). We then identified regions of variable chromatin accessibility (“peaks”) and variable gene expression across any of these four differentiation trajectories. This analysis revealed continuous trajectories of relevant gene expression and accessibility changes across developmental pseudotime in different regions of the intestine. For example, TMPRSS15, which encodes the protein that converts trypsinogen to trypsin in the duodenum, is highly expressed in the duodenum, where its expression gradually increases in more differentiated cells (Supplemental Figure 17). We next clustered these variable peaks and genes to identify sets with shared behavior (Figure 6B and 6C), revealing sets of peaks and genes that are open and expressed early in the differentiation pseudotime (e.g. in stem-cells) in all regions of the intestine, which we denote as Early (E). This cluster includes general markers of intestinal stem cells, including RGMB, SOX9, SMOC2, and LGR5, which are shared across the intestine. Other clusters of genes and peaks include those predominantly found in undifferentiated duodenum and jejunum (e.g. REG3A, SCTR), in differentiated small intestine cells (e.g. MTTP, APOA4, APOC3, MME), in undifferentiated colon (e.g. ROR2, PTGDR), and in differentiated colon (e.g. SCNN1B^51^, P3H2, MS4A12).

To identify both chromatin drivers of cluster specific regulation, as well as relevant cluster-specific gene expression programs, we computed TF motif enrichments in each cluster of peaks (Figure 6D) and KEGG pathway enrichment in each cluster of genes (Figure 6E). Groups of peaks accessible late in the differentiation trajectories were enriched for HNF4 and JUN/FOS motifs. As expected, genes primarily expressed late in the differentiation trajectory in the small intestine (Late Duodenum Jejunum cluster) were enriched for multiple metabolic KEGG pathways including fat digestion and absorption and protein digestion and absorption.

We next identified TFs whose gene expression is correlated with the activity of their motifs by computing correlation between TF expression and ChromVar deviation for the cells in each of the four differentiation trajectories. We found 68 TFs with expression correlated with their motif activity (r>0.7) and plotted the integrated gene expression of these transcription factors (Figure 6F). Many of these factors display similar activity along all four differentiation trajectories. For example, ASCL2, a master regulator in intestinal stem cells, is highly expressed at the beginning of all four trajectories. Other TFs, such as ETV6, exhibit different behaviors in different regions of the intestine. ETV6 is a transcription factor with decreased expression in colorectal cancer compared to normal colon, and genetic variation in ETV6 may confer colorectal cancer susceptibility ^52^. We found that ETV6 is more highly expressed in the colon than in the small intestine and, unlike the small intestine, ETV6 expression increases in more differentiated cells in the colon.

As an example of examining how genes with unique expression patterns may be controlled, we next tested which regulatory elements may be responsible for this variable expression of ETV6 in different regions of the intestine. We identified peaks with accessibility correlated to expression of ETV6 in any region of the intestine and then plotted how accessibility of these peaks changes along the differentiation pseudotime in each region (Figure 6G). For the seven most correlated peaks, accessibility tended to be highest in the colon, where expression is the greatest. The peak most accessible in the other regions of the intestine (Intronic 3) became less accessible along the differentiation trajectory, consistent with the decreased expression along the differentiation pseudotime in these regions. Similar behavior was also observed in the colon, where the Intronic 3 peak became less accessible along the differentiation trajectory despite expression of ETV6 increasing in the colon. However, multiple other peaks, both distal and intronic, exhibited increasing accessibility in more differentiated cells only in the colon, and we speculate that these regulatory elements may drive the increased expression of ETV6 in differentiated colon cells. The same logic can be applied to identify regulatory elements that may drive changes in gene expression throughout the intestine. For example, we identified 4 peaks that are highly correlated with expression of TMPRSS15 and may drive its increased expression in the duodenum (Supplemental Fig. 17). Taken together, this analysis provides a reference for the regulation of stem cell to enterocyte differentiation across the intestine.

We next tested if disease heritability is enriched in cell-type specific marker peaks in intestine cell-types using linkage disequilibrium (LD) score regression (Figure 6H, Methods). We identified a significant increase in heritability for Crohn’s disease and coeliac disease in T-cell marker peaks, consistent with the importance of T-cells in their pathogenesis^53^. We observed the most significant enrichment of heritability for BMI in enteroendocrine cells, suggesting that genetic variation may have an impact on enteroendocrine cells leading to effects on BMI. As a control, we also tested if heritability of GWAS SNPs was enriched in an unrelated trait (dental caries) and found no cell type specific enrichment. These results map important disease traits to specific cell types in the intestine.

## Discussion

Using a variety of different single cell (CODEX, snRNA-Seq, snATAC-Seq) technologies we analyzed many different regions of the intestine at high resolution. Our work greatly extends previous single cell studies by combining both spatial proteomic data sets (CODEX imaging) and snRNA and snATAC technologies. We demonstrated the extensive cellular complexity of the small intestine including considerable epithelial heterogeneity and novel secretory cell subtypes, that the different regions of the intestine have different cell compositions, and that cells are organized into different neighborhoods that also form communities. We also show conservation as well as heterogeneity of these neighborhoods and communities across the intestine. Finally, the open chromatin regulatory program was also mapped, defining key regulators and differentiation pathways utilized in the different regions of the intestine.

We generated a spatially hierarchical description of the intestine that is derived from the cell type labels we generate in our CODEX multiplexed imaging data. The first layer we computed was the immediate cellular neighborhoods within the tissues by examining conserved cell type composition within ten nearest neighbors of cells^10^. These represented and identified microstructures found within the intestine such as the vasculature or immune follicles. We further built upon the neighborhood concept to understand how neighborhoods of cells are interacting with each other and tend to colocalize. We quantitatively characterized these within communities of multicellular structures which can be further categorized into major community types. While the same *major communities* were shared between the small intestine and colon we observed differential localization of the *immune communities* localized to the apex in the colon and the base of the epithelium in the small intestine, likely reflective on differential microbial loads. This categorization isolated mucosal areas of the tissue enriched with immune and epithelial cells and defined overall structures and interactions of multicellular neighborhoods in comparing the colon and small intestine. This hierarchical view of multiplexed spatial data can serve as a template reference for other spatial atlasing efforts.

We focused our spatial analyses on immune cell organization within the intestine because plasma cells, CD4+ T cells, and CD8+ T cells all play critical roles in the intestinal immune response. Plasma cells play an important role in immunity by secreting IgA antibodies, the most abundantly produced antibody, critical for maintaining a homeostatic relationship with microbiota and food antigens^54, 55^. Interestingly, plasma cells had the highest same-cell density and were also found to co-localize with antigen-presenting cells in this multicellular neighborhood that was found in all areas of the intestine. By merging the snRNAseq data and CODEX data we also found that plasma cells and transit amplifying epithelial cells were one of the pairs of cells that co-localized more in the colon than the small intestine and differentially expressed SEMAD-MET ligand pair known to induce B cell clustering. Plasma cells require a special niche for survival in the bone marrow^56^ with survival factors such as CD44, a proliferation-inducing ligand (APRIL), IL-6, and SDF-1^56, 57^. Localization of plasma cells is critical in gut-associated lymphoid tissue (GALT) in the subepithelial space for transcytosis of IgA through the epithelium, and we found the *plasma cell enriched* neighborhood form an important crossroads of several other immune and epithelial neighborhoods within the mucosa. Consequently, our data suggests that antigen presenting cells and transit amplifying cells from the colon play critical roles in maintaining the rich plasma-cell zones of the mucosa that connect other multicellular immune and epithelial structures.

CD4+ T cells are involved both in CD4+ T cell support of B cell and CD8+ T cell activation. We found that CD4+ T cells were broadly distributed and enriched in all immune-rich multicellular structures (*innate immune enriched, inner follicle, outer follicle, adaptive immune enriched,* and *plasma cell enriched*) characteristic of their supportive functions. In particular, CD4+ T cells are the most enriched cell type in the *outer follicle* multicellular neighborhood, which was present in all sites of the intestine regardless of the presence of fully developed follicle structures. This observation suggests that the immune system appears structurally poised along the intestine to generate germinal center focused immune responses locally as needed. Follicle-type multicellular structures were among the least conserved structures when comparing the small intestine to the colon. Further understanding these differences will be important to clarify functional differences, development, and maintenance, as follicles are associated with beneficial cancer outcomes, increased autoimmunity, and clearance of infection^10, 58^. Moreover, the broad involvement of CD4+ T cells in diverse immune multicellular structures suggests their modulation might be akey therapeutic target for regulating immune responses in the intestine.

CD8+ T cells are critical for antiviral cytotoxic function and were one of the few cell types defined within the CODEX data shown to decrease from the small intestine to the colon. This compositional change likely reflects differences in antigen availability from both initial exposure as well as greater access to food antigens and foreign material with less mucus-secreting cells as compared to the colon^26, 59^. These differences were segregated into two neighborhoods *CD8+ T enriched IEL and adaptive immune*, where the *CD8+ T enriched IEL* neighborhood became nearly depleted in the colon. This structural difference further confirms the phenotypic distinction of IEL CD8+ T cells from the rest of CD8+ T cells within the intestine and merits additional study to understand maintenance, regulation, and renewal^60^ of these cells. Indeed, CD8+ T cells presence and density correlate with beneficial anti-cancer outcomes^61^ and cancer rates are increased within the colon as compared to the small intestine^62, 63^. It will be interesting to explore this correlation in the future and understand if intraepithelial immune cells help prune out defective epithelial cells.

We also calculated the similarity of multicellular structures from the small intestine to the colon and generated a conservation score to describe the compositional similarity. From this analysis it was apparent that all multicellular structures detected shared core features of the structure such as a high enrichment of CD4+ T cells for *outer follicle* structures, but varied on compositional enrichment of other cell types. The most varied structure was the *early secretory epithelial* multicellular neighborhood. This further substantiates the crypt structure as key regulators of differential function between the small intestine and colon^12^ and agrees with differences observed from RNA and ATAC datasets.

Leveraging paired transcriptome and chromatin accessibility data, we achieved further granularity to define the diversity of cell types in the intestine. Overall, different regions of the colon exhibited highly concordant cell type abundances. However, the cell type compositions of the small intestine regions were more diverse as compared to the colon, with the ileum often exhibiting immune and stromal cell type fractions shared between the small intestine regions and the colon. We observe greater diversity of specialized epithelial cells, including enteroendocrine and mucin-secreting cells in the small intestine. Within all regions, we identified goblet cells with high expression of MUC2. However, in the duodenum we identified two additional clusters containing cells characterized by high expression of the gel forming mucins, MUC5B and MUC6. The MUC6 cluster may represent cells of the duodenal Brunner’s glands, which have been shown to express high levels of MUC6^37^. The MUC5B cluster was more surprising as MUC5B is known to be expressed in goblet cells in the lung^64^ and gallbladder^65^, and at low levels in colon goblet cells^65^ but not in the duodenum, where MUC2 and MUC6 are the primary mucins.

While previous scRNA datasets have examined the diversity of cell types in the human intestine, our integrated snRNA and snATAC dataset provides a detailed single cell regulatory map of the intestine. We used this dataset to identify transcription factors in different cell types whose gene expression is highly correlated with the accessibility of the motif to which they bind. This identified TFs known to be master regulators of different cell types, including POU2F3 in Tuft cells, ASCL2 in stem cells, and RFX6 in enteroendocrine cells, while also nominating many additional transcription factors that are likely important regulators in their respective cell types. This includes RUNX1 and RUNX2 in Tuft cells, FOXA3 and ATOH1 in goblet cells, and ZBTB18 in Paneth cells.

Within all regions of the intestine, intestinal stem cells differentiate into mature absorptive enterocytes. By integrating gene expression and chromatin accessibility data along the differentiation trajectories in different regions of the intestine, we nominate TFs that exhibit consistent behavior across absorptive differentiation in all regions of the intestine. Among these are known intestinal stem cell regulators such as SOX9 and ASCL2, which are highly expressed and have high chromatin activity of their binding motif in cells at the beginning of the differentiation trajectory in all regions of the intestine. We also observe a number of differences in TF dynamics between the trajectories in different regions. One such example is ETV6, which is highly expressed in differentiated absorptive cells in the colon, but not other regions of the intestine. Interestingly, ETV6 expression is decreased in CRC and genetic variation in ETV6 may confer CRC risk^52^. Our data allow us to speculate on why this may be the case, as ETV6 is important for normal colon differentiation and thus loss of ETV6 may prevent differentiation of colon stem cells. Examining these data also allows us to link specific regulatory elements with the expression of transcription factors along the trajectory. For the case of ETV6, this analysis identifies one distal and three intronic regulatory elements with similar activity that are most likely responsible for driving expression of ETV6 along the absorptive differentiation trajectory in the colon.

Chromatin accessibility data also allow assessment of the cell-type specific enrichment with regulatory elements of common genetic variation linked to prevalent intestinal diseases (e.g. GWAS hits) such as Celiac, ulcerative colitis, and Crohn’s disease. This analysis can nominate the specific cell types through which these GWAS hits may be functioning, as well as the cell types that may be driving disease etiology. We find that T cells are most enriched in the autoimmune conditions, Celiac disease, ulcerative colitis, and Crohn’s. We also find that heritability of BMI is enriched in enteroendocrine cells, suggesting that variation in BMI is likely partially driven by genetic variation in regions that are functional in intestine enteroendocrine cells. We noted that hypertension is linked to endothelial cells which might represent a more general effect of endothelial cells; however, it would be interesting to explore if individuals with hypertension also have intestinal issues. Regardless, overall we can assign heritability of specific diseases to specific intestinal cell types.

There are several limitations to our study. First, for each patient, we typically analyzed a single sample from each intestinal region. Since the CODEX and the single cell data are largely in agreement and the variance across subjects is less than that across regions it is likely that these patterns are representative of these regions. It is also important to note that three of the four adults analyzed were older and three were male. The patterns across a wide range of ages and ethnicities remain to be elucidated. We are also underpowered to ascertain sex differences, which likely will be important given differences in disease risk for males and females^66^. These limitations can be addressed in the future with the acquisition of more samples and data.

In summary, we present a detailed map of the human intestine, and in particular the first multiplexed imaging reference for healthy small intestine and colon. In addition to biological insights this can serve as an important reference for intestinal diseases (e.g. IBD) as well as comparisons with other organisms.

## Methods

### Array Creation

Imaging data was collected from four human donors, each of whom constitutes a dataset. Each dataset includes two arrays of tissues that were imaged together on the same coverslip with four tissues per array: colon (sigmoid, descending, transverse, and ascending), and small intestine (ileum, mid-jejunum, proximal jejunum, duodenum). Arrays were constructed on the cryostat and sectioned at a width of 7 um.

### CODEX Antibody Conjugation and Panel Creation

CODEX multiplexed imaging was executed according to the CODEX staining and imaging protocol previously described^8^. Antibody panels were chosen to include targets that identify subtypes of intestinal epithelium and stromal cells, and cells of the innate and adaptive immune system. Detailed panel information can be found in supplementary table 5. Each antibody was conjugated to a unique oligonucleotide barcode, after which the tissues were stained with the antibody-oligonucleotide conjugates and validated that staining patterns matched expected patterns already established for IHC within positive control tissues of the intestine or tonsil. Similarly, Hematoxylin and Eosin morphology staining were used to confirm location of marker staining. First, antibody-oligonucleotide conjugates were tested in low-plex fluorescence assays and signal-to-noise ratio was also evaluated at this step, then they were tested all together in a single CODEX multicycle.

### CODEX Multiplexed Imaging

The tissue arrays were then stained with the complete validated panel of CODEX antibodies and imaged^8^. Briefly, this entails cyclic stripping, annealing, and imaging of fluorescently labeled oligonucleotides complementary to the oligonucleotide on the conjugate. After validation of the antibody-oligonucleotide conjugate panel, a test CODEX multiplexed assay was run, during which signal-to-noise ratio was again evaluated, and the optimal dilution, exposure time, and appropriate imagine cycle was evaluated for each conjugate (see supplementary table 5). Finally, each array underwent CODEX multiplexed imaging. Metadata from each CODEX run can be found in supplementary table 6.

### CODEX Data Processing

Raw imaging data were then processed using the CODEX Uploader for image stitching, drift compensation, deconvolution, and cycle concatenation. Processed data were then segmented using the CODEX Segmenter or CellVisionSegmenter, a watershed-based single-cell segmentation algorithm and a neural network R-CNN-based single-cell segmentation algorithm respectively. Both the CODEX Uploader and Segmenter software can be downloaded from our GitHub site (https://github.com/nolanlab/CODEX), and the CellVisionSegmenter software can be downloaded at https://github.com/bmyury/CellVisionSegmenter or https://github.com/michaellee1/CellSeg. After the upload, the images were again evaluated for specific signal: any markers that produced an untenable pattern or a low signal-to-noise ratio were excluded from the ensuing analysis. Uploaded images were visualized in ImageJ (https://imagej.nih.gov/ij/).

### Cell Type Analysis

B001 and B004 cell type identification were done following the methods developed previously^67^. Briefly, nucleated cells were selected by gating DRAQ5, Hoechst double-positive cells, followed by z-normalization of protein markers used for clustering (some phenotypic markers were not used in the unsupervised clustering). Then the data were overclustered with X-shift (https://github.com/nolanlab/vortex) or leiden-based clustering with the scanpy Python package. Clusters were assigned a cell type based on average cluster protein expression and location within image. Impure clusters were split or reclustered following mapping back to original fluorescent images.

### Cell Type Annotation using STELLAR

CODEX cell type labels were transferred to other donors (B005 & B006) using STELLAR as previously described (see companion *Nature Methods* manuscript submitted *Brbic et. al*). Briefly, STELLAR,a geometric deep learning method, utilized the spatial and molecular cell information to transfer cell type labels from B004 to normalized B005 and B006 datasets from other donors. Cell type labels were further refined with phenotypic markers such as M1 vs. M2 macrophages.

### CODEX Cell Density Analysis

CODEX same-cell density was analyzed by taking the average distance of the 5 nearest neighbors of the same cell type for each individual cell in each imaged region. This average distance was divided by the most diffuse distance for same cell types. The most diffuse same cell distance was calculated by taking the number of total cells of a given cell type divided by the total area of the region. Thus, numbers that are closer to one are least dense and numbers closer to 0 are more dense with cells of the same cell type.

### Neighborhood Identification Analysis

Neighborhood analysis was performed as described previously^10^. Briefly a window size of 10 nearest neighbors was taken across the tissue cell type maps and overclustered to 30 clusters. These clusters were mapped back to the tissue and evaluated for cell type enrichments to determine overall structure and merged down into 18 unique structures.

### Neighborhood Conservation Analysis

To determine neighborhood compositional (cell type) conservation across the small bowel and colon, neighborhood enrichment scores were found separately for both the small bowel and colon samples across all donors. This enrichment score is the average cell type percentage within the average of the neighborhood cluster divided by the average cell type percentage for all cells. The colon scores were subtracted from the small bowel scores to provide the heatmap that was ordered both in terms of greatest absolute sum of differences for both neighborhood and cell type in conservation.

### Community and Major Community Identification Analysis

Communities were determined similar to how multicellular neighborhoods were determined with some minor differences. Briefly, the cells in the neighborhood tissue maps were taken with a larger window size of 100 nearest neighbors. These windows were taken across the entirety of the tissue and the vectors then clustered with k-means clustering and overclustering with 20 total clusters. These clusters were mapped back to the tissue and evaluated for neighborhood composition and enrichment to determine overall community type. Communities were categorized into one of four major communities: epithelial, immune, stroma, or smooth muscle.

### Hierarchical Intestine Structural Graphs

Each hierarchical level was connected to the next by either what it contributed to the largest in the next level, or what made up at least 15% of this next hierarchy. The percentage of each feature in the level was represented by size of the shape. The shape and color combination correspond to the level and feature respectively. The size of the connecting line represents the amount a feature contributes to the next feature.

### Spatial Context Maps

Spatial context maps were created as previously described^34^. Briefly, first only cells which made up either epithelial or immune major cell types were selected for further analysis. Then, windows were taken again with the neighborhood types captured from the 100 nearest neighbors. Individual square combinations were selected by what combination of neighborhoods were needed to make up at least 85% of the total composition of the windows. Combinations were then counted and represented in size by the size of the black circle underneath the square neighborhood combination.

### Single-nuclei Experimental Methods: Tissue Dissociation and Nuclei Isolation

Nuclei were isolated using the OmniATAC protocol^68^. Isolation of nuclei was carried out on wet ice. 40-60mg of flash-frozen tissue was gently triturated and thawed in 1ml HB (Lysis) Buffer (1.0341x HB Stable solution, 1M DTT, 500 mM Spermidine, 150mM Spermine, 10% NP40, cOmplete Protease Inhibitor, Ribolock) for 5 minutes. Tissue was then dounced 10 times with pestle A and 20 times with pestle B, or until there was no resistance from either pestle. The samples were then filtered through a 40um cell strainer (Falcon; 352340) and the resulting homogenate was transferred to a pre-chilled 2ml LoBind tube. Samples were spun in a 4°C fixed angle centrifuge for 5 minutes at 350 RCF to pellet nuclei. After spinning, all but 50ul of supernatant was removed. 350ul HB was added to the nuclei pellet for a total volume of 400ul and nuclei were gently resuspended with a wide bore pipet. One volume of 50% Iodixanol (60% OptiPrep [Sigma Aldrich; D1556], Diluent Buffer [2M KCl, 1M MgCl2, 0.75M Tricine-KOH pH 7.8], Water) was added and the resulting solution was gently triturated. Next, 600ul of 30% Iodixanol was carefully layered under the 25% mixture. Finally, 600ul of 40% Iodixanol was layered under the 30% mixture. Sample was then spun for 20 min at 3,000 RCF in a 4°C swinging bucket centrifuge, resulting in a visible band of nuclei. Supernatant was aspirated down to within 200– 300ul of the nuclei band. The nuclei band was then collected at 200ul and transferred to a fresh 1.5ml tube. Sample was diluted with one volume (200ul) Resuspension Buffer (1x PBS, 1% BSA, 0.2u/uL Ribolock). Nuclei concentration was determined using the Countess II FL Automated Cell Counter (ThermoFisher; AMQAF1000).

### Single-nuclei Experimental Methods: Single-nuclei assay for transposase-accessible chromatin using sequencing (snATAC-seq)

snATAC-seq targeting 9,000 nuclei per sample was performed using Chromium Next GEM Single Cell ATAC Library & Gel Bead Kit v1.1 (10x Genomics, 1000175) and Chromium Next GEM Chip H (10x Genomics, 1000161) or Chromium Single Cell ATAC Library & Gel Bead Kit (10x Genomics, 1000110). Libraries were sequenced on an Illumina NovaSeq 6000 (1.4 pM loading concentration, 50 × 8 × 16 × 49 bp read configuration) targeting an average of 25,000 reads per nuclei.

### Single-nuclei Experimental Methods: Single-nuclei transcriptome sequencing (snRNA-seq)

snRNA-seq targeting 9,000 nuclei per sample was performed using Chromium Next GEM Single Cell 3’ Reagent Kits v3.1 (10x Genomics, 1000121) and Chromium Next GEM Chip G Single Cell Kit (10x Genomics, 1000120). Libraries were pooled and sequenced on an Illumina NovaSeq 6000 (Read 1 = 28bp, i7 index=8bp, i5 index=0bp, Read 2=91bp read configuration) targeting an average of 20,000 reads per nuclei.

### Single-nuclei Experimental Methods: Single-nuclei multiome experiments

snMultiome experiments targeting 9,000 nuclei per sample were performed using Chromium Chromium Next GEM Single Cell Multiome ATAC + Gene Expression (10x Genomics, 1000283). ATAC (Read 1 = 50bp, i7 index=8bp, i5 index=24bp, Read 2=49bp read configuration) and RNA (Read 1 = 28bp, i7 index=10bp, i5 index=10bp, Read 2=90bp read configuration) libraries were sequenced separately on an Illumina NovaSeq 6000.

### Single-nuclei Analytical Methods: Initial processing of single-nuclei data

Initial processing of scATAC-seq data was performed using the Cell Ranger ATAC Pipeline (go.10xgenomics.com/scATAC/cell-ranger-ATAC) by first running cellranger-atac mkfastq to demultiplex the bcl files and then running cellranger-atac count to generate scATAC fragments files. Initial processing of snRNA-seq data was done with the Cell Ranger Pipeline (https://support.10xgenomics.com/single-cell-gene-expression/software/pipelines/latest/what-is-cell-ranger) by first running cellranger mkfastq to demultiplex the bcl files and then running cellranger count. Since nuclear RNA was sequenced, data were aligned to a pre-mRNA reference. Initial processing of the mutiome data, including alignment and generation of fragments files and expression matrices, was performed with the Cell Ranger ARC Pipeline.

### Colocalization Analyses

The CODEX data was used to compute and compare the colocalization quotient (CLQ) between all cell-type pairs in the small bowel versus the colon. The colocalization quotient between cell type A and cell type B was calculated using the expression 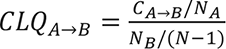, where CADB is the number of cells of cell type A among the defined nearest neighbors of cell type B. N is the total number of cells and NA and NB are the numbers of cells for cell type A and cell type B. The student’s t-test, adjusted for multiple hypothesis testing, was to identity statistically significant different CLQs between small bowel and colon.

### Ligand and Receptor Analyses

The FANTOM5 database^70^ and 12 more literature-supported experimentally-validated ligand and receptor pairs were used to obtain the final list of validated ligand receptor pairs (Yu et al., 2021).

Non-parametric Wilcoxon rank-sum test was used to identify differentially expressed ligands and receptors in the small bowel versus the colon (adjusted p-value cutoff = 0.05).

### Single-nuclei Analytical Methods: Quality control, dimensionality reduction, and clustering of snATAC data

The snATAC fragments files were loaded into R (v4.0.2) using the createArrowFiles function in ArchR^43^. Quality control metrics were computed for each cell and any cells with TSS enrichments less than 4 were removed. Cells were also filtered based on the number of unique fragments sequenced using a unique fragment cutoff that was defined for each individual sample. The sample-specific cutoffs enabled us to account for differences in sequencing depth between samples and ranged from 1,000 to 5,000, with the most common cutoff being 3000 fragments/cell. After quality control and filtering, doublet scores were computed with the ArchR function addDoubletScores with k=10, knnMethod = “UMAP”, and LSIMethod = 1. An ArchR project was then created and doublets were filtered with filterDoublets with a filter ratio of 1.2. An IterativeLSI dimensionality reduction was then generated on all cells using addIterativeLSI, with iterations = 3 and varFeatures = 15000. Next, clusters were added with addClusters with resolution = 2 and the resulting clusters were divided into groups based on if the cells exhibited high gene activity scores^43^, a measure of accessibility within and around a gene body, for known immune, stromal, or epithelial marker genes.

### Single-nuclei Analytical Methods: Quality control, dimensionality reduction, and clustering of snRNA data

After running Cell Ranger, the filtered_feature_bc_matrix produced by Cell Ranger was read into R with the Seurat^71^ function Read10X. The data was filtered to remove nuclei with fewer than 400 unique genes per nuclei or greater than 4000 genes per nuclei. DoubletFinder^72^ was run for each sample using PCs 1-20. nExp was set to 0.08*nCells^2^/10000, pN to 0.25, and pK to 0.09, and cells classified as doublets were removed prior to downstream analysis.

The remaining cells from all samples were merged into a single seurat object, and nuclei with greater than 10,000 counts/nuclei or greater than 10% mitochondrial RNA were removed. The data was then processed with Seurat’s standard pipeline^71^. First, NormalizeData was run using the method LogNormalize and scale.factor of 10,000. Variable features were identified with Seurat’s findVariableFeatures using the vst method and 2,000 features. ScaleData was then run on all genes and PCs were computed with RunPCA. To account for batch effects between different donors in clustering, the RunHarmony^73^ function was run with the data being grouped by Donor (e.g. B001, B004, B005, and B006). RunUMAP was then used to generate a umap dimensionality reduction from the harmony reduction and the cells were clustered by first using FindNeighbors with reduction = “harmony” and dims = 1:20 and then FindClusters with a resolution of 1. Expression of marker genes in the resulting clusters were then used to label clusters as epithelial, stromal, or immune for downstream analysis.

### Single-nuclei Analytical Methods: Annotation of single-nuclei data

The snATAC and snRNA data were annotated in the following groups: epithelial duodenum, epithelial jejunum, epithelial ileum, epithelial colon, stromal, and immune. For the ATAC data, the cells in each of these compartments were subset into a new ArchR project. addIterativeLSI was then run for each compartment. addHarmony was then run using the LSI dimensions as input. Following computation of the harmony dimensions, the cells were clustered using addClusters and a UMAP was computed based on the harmony coordinates with addUMAP. Clusters were annotated by examining gene activity scores of known marker genes. Marker genes were used for initial annotation of cell types including Best4+ Enterocytes (BEST4, OTOP2), Goblet (MUC2, TFF1, SYTL2), Immature Goblet (KLK1, RETNLB, CLCA1), Stem (RGMB, SMOC2, LGR5, ASCL2), Tuft (SH2D6, TRPM5, BMX, LRMP, HCK), Enteroendocrine (SCGN, FEV, CHGA, PYY, GCG), CyclingTA (TICRR, CDC25C), Paneth (LYZ, DEFA5), CD4+ and CD8+ T-cells (CD2, CD3E, IL7R, CD4, CD8), B-cells (PAX5, MS4A1, CD19), Plasma cells (IGLL5, AMPD1), NK cells

(SH2D1B), macrophages/monocytes (CD14), and mast cells (HDC, GATA2, TPSAB1). Smooth muscle/myofibroblast clusters exhibited high expression of ACTA2, MTH11, and TAGLN. Villus fibroblasts exhibited high expression of WNT5B and some crypt fibroblasts exhibited high expression of RSPO3. For the snRNA cells, cells were divided into 6 Seurat objects from the compartments listed above. Cells from each compartment were run through the same pipeline listed above (NormalizeData, ScaleData, RunHarmony, FindNeighbors, and FindClusters). For the immune and stromal compartments, we also labeled the scATAC datasets with previously published scRNA datasets using addGeneIntegrationMatrix, and the annotations were generally consistent with the manual annotations. Notably the smooth muscle/myofibroblast 1 cluster separated into multiple clusters in RNA space. In annotating these clusters, we labeled all clusters as smooth muscle/myofibroblast 1 to match the snATAC data, but they could have been divided into additional clusters. We also found that some nuclei that clustered with epithelial cells did not have clear expression of marker genes and generally were slightly lower quality in terms of genes/nuclei. We labeled these cells as “Epithelial” but did not give them more detailed annotations. We also note that a small number of nuclei that were thought to be doublet clusters were also removed during annotation. Following initial annotation of epithelial cells, enteroendocrine cells in the snRNA data were subclustered, and known subtypes of enteroendocrine cells were annotated based on expression of marker genes (Figure 5C).

### Single-nuclei Analytical Methods: Integration of snRNA and snATAC data and nomination of regulatory TFs

The snRNA and snATAC datasets from the four primary regions of the intestine (duodenum, jejunum, ileum, and colon) were integrated separately using the ArchR function addGeneIntegrationMatrix with reducedDims = “Harmony” and useMatrix = “GeneScoreMatrix”. We then identified TF regulators following the ArchR manual for identifying TF regulators for each region, with a correlation cutoff of 0.6. TFs that met the criteria for regulators in any of the four primary regions of the intestine are plotted in Figure 5.

### Single-nuclei Analytical Methods: Annotation of single-nuclei data: Analysis of absorptive differentiation trajectories

Absorptive differentiation trajectories for each main section of the colon (duodenum, jejunum, ileum, and colon) were inferred by running the ArchR function addTrajectory with trajectory = c(“Stem”, “TA2”, “TA1”, “Immature Enterocytes”, “Enterocytes”), reducedDims set to the harmony dimensions, and groupBy set to “CellType.” To identify variable peaks along the trajectory, a matrix of accessibility in all peaks along the trajectory was first generated with getTrajectory with useMatrix = “PeakMatrix” and log2Norm = TRUE. Peaks with variance>0.9 in any of the four regions were then identified with the function plotTrajectoryHeatmap with varCutOff = 0.9, returnMatrix = TRUE, scaleRows = FALSE, and maxFeatures = 100000. The four matrices returned by getTrajectory were then concatenated into a single matrix and the matrix was subset to include only peaks that met the variance criteria of 0.9 in at least one of the four regions and had an absolute difference in magnitude of at least 0.2. Row z-scores for the resulting matrix were computed with the ArchR function .rowZscore. The resulting row z-scores were kmeans clustered using the function kmeans with the number of clusters set to 7 and iter.max = 500. One cluster of peaks did not show a characteristic pattern and was not included in Figure 6. Hypergeometric enrichment of motifs in marker peaks was computed with peakAnnoEnrichment and the resulting p values are plotted in Figure 6D. Variable genes along the trajectory were identified with an analogous method, using GeneIntegrationMatrix instead of PeakMatrix when running getTrajectory and plotTrajectoryHeatmap, and the row z-scores were again kmeans clustered using the function kmeans with the number of clusters set to 7 and iter.max = 500. Enrichment of kegg pathways in these clusters of genes was determined with the limma function kegga^74^, and the resulting unadjusted p values are plotted in Figure 6E. Plots of integrated gene expression versus pseudotime were generated with the ArchR function plotTrajectory with default parameters. TFs with correlated motif activity and RNA expression were identified with correlateTrajectories as outlined in the ArchR manual. TFs that were correlated with expression in any of the four trajectories were included in the heatmap in Figure 6F. Plots of integrated gene expression along each trajectory were generated with plotTrajectory. Peaks correlated with gene expression were identified with addPeak2GeneLinks. To identify possible peaks linked to genes changing along the differentiation trajectory. In Figure 6G, the set of peaks that were correlated with ETV6 expression with a correlation of at least 0.65 in one of the four main intestinal regions was determined. The smoothed trajectory peak accessibility for each of these peaks was then plotted along the differentiation trajectory.

### Cell-type specific linkage disequilibrium (LD) score regression

To run cell type specific LD score regression, we first computed marker peaks for course cell types in our dataset. To do this we added cell type annotations to the full ArchR project will all cells and then defined a peak set for this object by running addGroupCoverages with groupBy = “CellType” followed by addReproduciblePeakSet and addPeakMatrix. We then defined less granular cell types by merging all myofibroblast clusters and pericytes into a single group, all fibroblast clusters into a single group, all non-stem absorptive epithelial cells into a single group, all enteroendocrine cells into a single group, CD4+ and CD8+ T cells into a single group, and lymphatic endothelial and endothelial cells into a single group. We determined marker peaks for the resulting groups of cells with getMarkerFeatures and ten selected peaks with getMarkers with cutOff = “FDR <= 0.1 & Log2FC >= 0.5”. The resulting peaks were then lifted over to hg19 from hg38. We then followed the LD score regression tutorial (https://github.com/bulik/ldsc/wiki) for cell-type specific analysis^75–77^. We used summary statistics from a number of UKBB traits (http://www.nealelab.is/uk-biobank/) related to the intestine, including Non-cancer illness code;self-reported: diverticular disease/diverticulitis, Non-cancer illness code;self-reported: crohns disease, Non-cancer illness code;self-reported: ulcerative colitis, Non-cancer illness code;self-reported: malabsorption/coeliac disease, Body mass index (BMI) as well as traits less clearly related to the intestine including Non-cancer illness code;self-reported: hypertension and Diagnoses - main ICD10: K02 Dental caries. Coefficient p-values from ldsc are plotted in the heatmap in Figure 6. Significance was determined by correcting the coefficient p-values for the number of cell-types tested with bonferroni correction with the R function p.adjust.

### Data Availability Statement

All of the published datasets in this study can be visualized and assessed through a website portal (https://portal.hubmapconsortium.org/).

## Acknowledgments

We thank the patients for donating their tissues. This work was supported by the U.S. National Institutes of Health (3U54HG010426); Cancer Research UK (C27165/A29073); the Parker Institute for Cancer Immunotherapy (PICI0025); Hope Realized Medical Foundation (209477). J.W.H. was supported by an NIH T32 Fellowship (T32CA196585) and an American Cancer Society—Roaring Fork Valley Postdoctoral Fellowship (PF-20-032-01-CSM). Figures 1A and 2A were created with Biorender.com.

## Author contributions

S.B. and C.M.S. developed the original CODEX panel and acquired the CODEX data from B001. V.V. helped establish CODEX data processing pipelines and integration with HuBMAP data transfers. M.B., K.C., J.W.H., and J.L. created STELLAR that was used for cell type label transfer of the CODEX data. J.W.H., S.B., and C.C. developed gut-specific CODEX antibodies and ran remaining donor samples. J.W.H. processed and analyzed CODEX multiplexed imaging data. R.C., D.C., S.A.N., A.M.H., and W.R.B. established experimental protocols and collected the snRNA and snATAC data. A.E.P. analyzed CODEX and snRNAseq data for neighborhood crosstalk. W.R.B. analyzed snATAC and snRNA data. W.R.B., J.W.H., W.J.G., and M.S. wrote the manuscript and created figures. D.C. handled data management and submission to repositories. A.H.and A.K.W were involved in project coordination. T.L. assessed morphological staining of samples. E.E. assisted in metadata annotation and biological interpretation. Y.L., Z.Z. and S.L. procured samples. Y.L., S.K.P., G.P. N., W.J.G. and M.S. supervised the study. All authors revised the manuscript and accepted its final version.

