## Supplemental Table 1 for "High Resolution Single Cell Maps Reveals Distinct Cell Organization and Function Across Different Regions of the Human Intestine"

Supplementary Table 1. Donor specific metadata.

| Stanford ID | B001 | B004 | B005 | B006 |
| --- | --- | --- | --- | --- |
| donor age | 67 | 78 | 24 | 38 |
| donor sex | female | male | female | male |
| donor race | White | Black | White | White |
| BMI | 30.2 | 35.1 | 23.2 | 29 |
| History of diabetes | yes | yes | no | no |
| History of cancer | no | no | no | no |
| History of hypertension | yes | yes | no | no |
| History gastrointestinal disease | no | no | no | no |
