## Supplemental Table 2 for "High Resolution Single Cell Maps Reveals Distinct Cell Organization and Function Across Different Regions of the Human Intestine"

Supplementary table 2. Relationship between snRNAseq and CODEX cell type annotations

| **Cell Transcriptomics** | **Cell CODEX** |
| --- | --- |
| B Cells | B |
| CD4+ | CD4+ T cell |
| CD8+ | CD7+ Immune |
| CD8+ | CD8+ T |
| Cycling TA | Cycling TA |
| Stem | Cycling TA |
| Stem2 | Cycling TA |
| + | DC |
| Endothelial | Endothelial |
| Best4+Enterocytes | Enterocyte |
| Enterocytes | Enterocyte |
| Tuft | Enterocyte |
| Goblet | Goblet |
| Secretory unknown 1 | Goblet |
| Secretory unknown 2 | Goblet |
| Immature Goblet | ITLN+ Enterocyte |
| Lymphatic endothelial cells | Lymphatic |
| Mono_Macrophages | M1 Macrophage |
| Mono_Macrophages | M2 Macrophage |
| Glia | Nerve |
| Neurons | Nerve |
| Enteroendocrine1 | Neuroendocrine |
| Enteroendocrine2 | Neuroendocrine |
| Enteroendocrine3 | Neuroendocrine |
| Epithelial | * |
| + | Neutrophil |
| NK | NK |
| Paneth | Paneth |
| Plasma | Plasma |
| Myofibroblasts 1 | Smooth muscle |
| Myofibroblasts 2 | Smooth muscle |
| Myofibroblasts 3 | Smooth muscle |
| Pericycles | Smooth muscle |
| Crypt Fibroblasts 1 | Stroma |
| Crypt Fibroblasts 2 | Stroma |
| Crypt Fibroblasts 3 | Stroma |
| Crypt Fibroblasts RSPO3+ | Stroma |
| Villus Fibroblasts WNT5B+ | Stroma |
| TA1 | TA |
| TA2 | TA |
| Mast | x |

* Class not well defined; + cell type not identified in snRNA;
 + cell type not identified in CODEX
