## Supplemental Table 3 for "High Resolution Single Cell Maps Reveals Distinct Cell Organization and Function Across Different Regions of the Human Intestine"

Supplementary table 3. Uniqueness of ligand-receptor pairs across colocalized cells in large colon

| Pair of ligand-receptor | Pair of cells | N times appears | % across all pairs of cell types |
| --- | --- | --- | --- |
| HLA-C_LILRB1 | Enterocytes-Plasma | 1 | 0 |
| HLA-B_LILRB1 | Enterocytes-Plasma | 2 | 0.15 |
| HLA-B_LILRB1 | TA1-Plasma | 2 | 0.15 |
| PSEN1_NOTCH3 | Enterocytes-Pericytes | 3 | 0.3 |
| PSEN1_NOTCH3 | Immature Enterocytes-Pericytes | 3 | 0.3 |
| FGF2_SDC4 | Myofibroblasts 1-Enterocytes | 4 | 0.46 |
| SEMA4D_PLXNB1 | Plasma-Best4 Enterocytes | 4 | 0.46 |
| JAG1_NOTCH3 | Immature Enterocytes-Pericytes | 6 | 0.76 |
| JAG1_NOTCH3 | Tuft-Pericytes | 6 | 0.76 |
| IGFBP4_LRP6 | Pericytes-Immature Enterocytes | 6 | 0.76 |
| MLLT4_F11R | Myofibroblasts 1-Enterocytes | 7 | 0.91 |
| EFNB2_EPHB1 | Best4 Enterocytes-Myofibroblasts 1 | 8 | 1.06 |
| EFNB2_EPHB1 | Enterocytes-Myofibroblasts 1 | 8 | 1.06 |
| BMP2_BMPR1B | Enterocytes-Myofibroblasts 3 | 8 | 1.06 |
| EFNB2_EPHB1 | Immature Enterocytes-Myofibroblasts 1 | 8 | 1.06 |
| TNC_PTPRB | Myofibroblasts 1-Best4 Enterocytes | 12 | 1.67 |
| TNC_PTPRB | Myofibroblasts 1-Enterocytes | 12 | 1.67 |
| TNC_PTPRB | Myofibroblasts 1-Immature Enterocytes | 12 | 1.67 |
| FN1_ITGA3 | Plasma-TA1 | 13 | 1.82 |
| EFNA5_EPHB1 | Best4 Enterocytes-Myofibroblasts 1 | 14 | 1.97 |
| HSPG2_PTPRS | Best4 Enterocytes-Myofibroblasts 1 | 14 | 1.97 |
| BMP2_BMPR1A | Enterocytes-Myofibroblasts 1 | 14 | 1.97 |
| BMP2_BMPR2 | Enterocytes-Myofibroblasts 1 | 14 | 1.97 |
| EFNA5_EPHB1 | Enterocytes-Myofibroblasts 1 | 14 | 1.97 |
| HSPG2_PTPRS | Enterocytes-Myofibroblasts 1 | 14 | 1.97 |
| BMP2_BMPR1A | Enterocytes-Myofibroblasts 3 | 14 | 1.97 |
| EFNA5_EPHB1 | Immature Enterocytes-Myofibroblasts 1 | 14 | 1.97 |
| HSPG2_PTPRS | Immature Enterocytes-Myofibroblasts 1 | 14 | 1.97 |
| EFNA5_EPHB1 | Tuft-Myofibroblasts 1 | 14 | 1.97 |
| ADAM17_ITGA5 | Enterocytes-Myofibroblasts 1 | 15 | 2.12 |
| ADAM17_ITGA5 | Immature Enterocytes-Myofibroblasts 1 | 15 | 2.12 |
| EFNB2_EPHA6 | Best4 Enterocytes-Myofibroblasts 1 | 16 | 2.28 |
| EFNB2_EPHA6 | Best4 Enterocytes-Myofibroblasts 3 | 16 | 2.28 |
| EFNB2_EPHA6 | Enterocytes-Myofibroblasts 1 | 16 | 2.28 |
| EFNB2_EPHA6 | Enterocytes-Myofibroblasts 3 | 16 | 2.28 |
| EFNB2_EPHA6 | Immature Enterocytes-Myofibroblasts 1 | 16 | 2.28 |
| EFNB2_EPHA6 | Immature Enterocytes-Myofibroblasts 3 | 16 | 2.28 |
| FGF2_FGFR2 | Myofibroblasts 1-Enterocytes | 16 | 2.28 |
| FGF2_FGFR2 | Myofibroblasts 1-Immature Enterocytes | 16 | 2.28 |
| FGF2_CD44 | Myofibroblasts 1-Tuft | 16 | 2.28 |
| FGF2_FGFR2 | Myofibroblasts 1-Tuft | 16 | 2.28 |
| LAMB1_ITGA6 | Best4 Enterocytes-Plasma | 16 | 2.28 |
| LAMC2_ITGA6 | Best4 Enterocytes-Plasma | 16 | 2.28 |
| LAMC2_ITGA6 | Enterocytes-Plasma | 16 | 2.28 |
| LAMC2_ITGA6 | Immature Enterocytes-Plasma | 16 | 2.28 |
| LAMB1_ITGA6 | TA2-Plasma | 16 | 2.28 |
| NLGN1_NRXN3 | Myofibroblasts 1-Best4 Enterocytes | 18 | 2.58 |
| NLGN1_NRXN3 | Myofibroblasts 3-Best4 Enterocytes | 18 | 2.58 |
| NLGN1_NRXN3 | Pericytes-Best4 Enterocytes | 18 | 2.58 |
| NLGN1_NRXN3 | Myofibroblasts 1-Tuft | 18 | 2.58 |
| NLGN1_NRXN3 | Myofibroblasts 3-Tuft | 18 | 2.58 |
| NLGN1_NRXN3 | Pericytes-Tuft | 18 | 2.58 |
| CDH1_PTPRM | Enterocytes-Myofibroblasts 1 | 20 | 2.88 |
| CDH1_PTPRM | Immature Enterocytes-Myofibroblasts 1 | 20 | 2.88 |
| ADAM9_ITGA6 | Enterocytes-Plasma | 20 | 2.88 |
| ADAM9_ITGA6 | Immature Enterocytes-Plasma | 20 | 2.88 |
| MLLT4_EPHB2 | Myofibroblasts 1-Best4 Enterocytes | 21 | 3.03 |
| MLLT4_EPHB2 | Myofibroblasts 1-Tuft | 21 | 3.03 |
| COL1A1_CD44 | Myofibroblasts 1-Tuft | 24 | 3.49 |
| FGF13_SCN8A | Myofibroblasts 3-Best4 Enterocytes | 25 | 3.64 |
| FN1_ITGB6 | Myofibroblasts 1-Enterocytes | 26 | 3.79 |
| FN1_ITGB6 | Myofibroblasts 2-Enterocytes | 26 | 3.79 |
| FN1_ITGB6 | Myofibroblasts 3-Enterocytes | 26 | 3.79 |
| FN1_ITGB6 | Pericytes-Enterocytes | 26 | 3.79 |
| FN1_ITGB6 | Myofibroblasts 1-Immature Enterocytes | 26 | 3.79 |
| FN1_ITGB6 | Myofibroblasts 2-Immature Enterocytes | 26 | 3.79 |
| FN1_ITGB6 | Myofibroblasts 3-Immature Enterocytes | 26 | 3.79 |
| FN1_ITGB6 | Pericytes-Immature Enterocytes | 26 | 3.79 |
| FN1_ITGB6 | Plasma-Enterocytes | 26 | 3.79 |
| FN1_ITGB6 | Plasma-Immature Enterocytes | 26 | 3.79 |
| THBS1_ITGA4 | Enterocytes-Plasma | 27 | 3.95 |
| THBS1_ITGA4 | Immature Enterocytes-Plasma | 27 | 3.95 |
| THBS1_ITGA4 | TA1-Plasma | 27 | 3.95 |
| THBS1_ITGA4 | TA2-Plasma | 27 | 3.95 |
| EFNA5_EPHA6 | Best4 Enterocytes-Myofibroblasts 1 | 28 | 4.1 |
| EFNA5_EPHA6 | Best4 Enterocytes-Myofibroblasts 3 | 28 | 4.1 |
| EFNA5_EPHA6 | Enterocytes-Myofibroblasts 1 | 28 | 4.1 |
| EFNA5_EPHA6 | Enterocytes-Myofibroblasts 3 | 28 | 4.1 |
| EFNA5_EPHA6 | Immature Enterocytes-Myofibroblasts 1 | 28 | 4.1 |
| EFNA5_EPHA6 | Immature Enterocytes-Myofibroblasts 3 | 28 | 4.1 |
| EFNA5_EPHA6 | Tuft-Myofibroblasts 1 | 28 | 4.1 |
| EFNA5_EPHA6 | Tuft-Myofibroblasts 3 | 28 | 4.1 |
| EFNA5_EPHA2 | Myofibroblasts 1-Enterocytes | 28 | 4.1 |
| EFNA5_EPHA2 | Myofibroblasts 1-Immature Enterocytes | 28 | 4.1 |
| SEMA4D_MET | Plasma-Enterocytes | 28 | 4.1 |
| SEMA4D_MET | Plasma-Immature Enterocytes | 28 | 4.1 |
| SEMA4D_MET | Plasma-TA2 | 28 | 4.1 |
| COL1A2_CD44 | Myofibroblasts 1-Tuft | 32 | 4.7 |
| TNC_EGFR | Myofibroblasts 1-Best4 Enterocytes | 36 | 5.31 |
| TNC_EGFR | Myofibroblasts 1-Enterocytes | 36 | 5.31 |
| TNC_EGFR | Myofibroblasts 1-Immature Enterocytes | 36 | 5.31 |
| THBS1_ITGA6 | Enterocytes-Plasma | 36 | 5.31 |
| THBS1_ITGA6 | Immature Enterocytes-Plasma | 36 | 5.31 |
| THBS1_ITGA6 | TA1-Plasma | 36 | 5.31 |
| THBS1_ITGA6 | TA2-Plasma | 36 | 5.31 |
| IL18_IL1RAPL1 | Best4 Enterocytes-Myofibroblasts 1 | 40 | 5.92 |
| IL18_IL1RAPL1 | Best4 Enterocytes-Myofibroblasts 2 | 40 | 5.92 |
| IL18_IL1RAPL1 | Enterocytes-Myofibroblasts 1 | 40 | 5.92 |
| IL18_IL1RAPL1 | Enterocytes-Myofibroblasts 2 | 40 | 5.92 |
| IL18_IL1RAPL1 | Immature Enterocytes-Myofibroblasts 1 | 40 | 5.92 |
| IL18_IL1RAPL1 | Immature Enterocytes-Myofibroblasts 2 | 40 | 5.92 |
| IL18_IL1RAPL1 | Tuft-Myofibroblasts 1 | 40 | 5.92 |
| IL18_IL1RAPL1 | Tuft-Myofibroblasts 2 | 40 | 5.92 |
| FGF7_FGFR2 | Myofibroblasts 1-Enterocytes | 40 | 5.92 |
| FGF7_FGFR2 | Myofibroblasts 3-Enterocytes | 40 | 5.92 |
| FGF7_FGFR2 | Myofibroblasts 1-Immature Enterocytes | 40 | 5.92 |
| FGF7_FGFR2 | Myofibroblasts 3-Immature Enterocytes | 40 | 5.92 |
| FGF7_FGFR2 | Myofibroblasts 1-Tuft | 40 | 5.92 |
| FGF7_FGFR2 | Myofibroblasts 3-Tuft | 40 | 5.92 |
| EFNA5_EPHB2 | Myofibroblasts 1-Best4 Enterocytes | 42 | 6.22 |
| MLLT4_NRXN3 | Myofibroblasts 1-Best4 Enterocytes | 42 | 6.22 |
| EFNA5_EPHB2 | Myofibroblasts 1-Tuft | 42 | 6.22 |
| MLLT4_NRXN3 | Myofibroblasts 1-Tuft | 42 | 6.22 |
| ANXA1_EGFR | Myofibroblasts 1-Best4 Enterocytes | 45 | 6.68 |
| ANXA1_EGFR | Myofibroblasts 3-Best4 Enterocytes | 45 | 6.68 |
| ANXA1_EGFR | Myofibroblasts 1-Enterocytes | 45 | 6.68 |
| ANXA1_EGFR | Myofibroblasts 3-Enterocytes | 45 | 6.68 |
| ANXA1_EGFR | Myofibroblasts 1-Immature Enterocytes | 45 | 6.68 |
| ANXA1_EGFR | Myofibroblasts 3-Immature Enterocytes | 45 | 6.68 |
| CDH1_IGF1R | Enterocytes-Myofibroblasts 1 | 48 | 7.13 |
| CDH1_IGF1R | Immature Enterocytes-Myofibroblasts 1 | 48 | 7.13 |
| FN1_PLAUR | Myofibroblasts 1-Enterocytes | 52 | 7.74 |
| FN1_PLAUR | Myofibroblasts 2-Enterocytes | 52 | 7.74 |
| FN1_PLAUR | Myofibroblasts 3-Enterocytes | 52 | 7.74 |
| FN1_PLAUR | Pericytes-Enterocytes | 52 | 7.74 |
| FN1_PLAUR | Myofibroblasts 1-Immature Enterocytes | 52 | 7.74 |
| FN1_PLAUR | Myofibroblasts 2-Immature Enterocytes | 52 | 7.74 |
| FN1_PLAUR | Myofibroblasts 3-Immature Enterocytes | 52 | 7.74 |
| FN1_PLAUR | Pericytes-Immature Enterocytes | 52 | 7.74 |
| FN1_PLAUR | Plasma-Enterocytes | 52 | 7.74 |
| FN1_PLAUR | Plasma-Immature Enterocytes | 52 | 7.74 |
| VIM_CD44 | Pericytes-Tuft | 56 | 8.35 |
| SEMA3C_NRP2 | Best4 Enterocytes-Myofibroblasts 1 | 60 | 8.95 |
| SEMA3C_NRP2 | Best4 Enterocytes-Myofibroblasts 3 | 60 | 8.95 |
| SEMA3C_NRP2 | Enterocytes-Myofibroblasts 1 | 60 | 8.95 |
| SEMA3C_NRP2 | Enterocytes-Myofibroblasts 3 | 60 | 8.95 |
| SEMA3C_NRP2 | Immature Enterocytes-Myofibroblasts 1 | 60 | 8.95 |
| SEMA3C_NRP2 | Immature Enterocytes-Myofibroblasts 3 | 60 | 8.95 |
| SEMA3C_NRP2 | Tuft-Myofibroblasts 1 | 60 | 8.95 |
| SEMA3C_NRP2 | Tuft-Myofibroblasts 3 | 60 | 8.95 |
| CNTN4_PTPRG | Best4 Enterocytes-Myofibroblasts 1 | 99 | 14.87 |
| CNTN4_PTPRG | Best4 Enterocytes-Myofibroblasts 2 | 99 | 14.87 |
| CNTN4_PTPRG | Immature Enterocytes-Myofibroblasts 1 | 99 | 14.87 |
| CNTN4_PTPRG | Immature Enterocytes-Myofibroblasts 2 | 99 | 14.87 |
| CNTN4_PTPRG | Tuft-Myofibroblasts 1 | 99 | 14.87 |
| CNTN4_PTPRG | Tuft-Myofibroblasts 2 | 99 | 14.87 |
| FN1_CD44 | Myofibroblasts 1-Tuft | 104 | 15.63 |
| FN1_CD44 | Myofibroblasts 2-Tuft | 104 | 15.63 |
| FN1_CD44 | Myofibroblasts 3-Tuft | 104 | 15.63 |
| FN1_CD44 | Pericytes-Tuft | 104 | 15.63 |
| FN1_CD44 | Plasma-Tuft | 104 | 15.63 |
| EFNA5_EPHA4 | Myofibroblasts 1-Best4 Enterocytes | 112 | 16.84 |
