## Supplemental Table 4 for "High Resolution Single Cell Maps Reveals Distinct Cell Organization and Function Across Different Regions of the Human Intestine"

Supplementary table 4: KEGG significant pathways from upregulated genes in colon associated with MET receptor

| ID | Description | GeneRatio | p-value | p.adjust | Count |
| --- | --- | --- | --- | --- | --- |
| hsa04520 | Adherens junction | 14/421 | 1.3E-05 | 2.4E-03 | 14 |
| hsa04360 | Axon guidance | 23/421 | 6.4E-05 | 6.3E-03 | 23 |
| hsa04015 | Rap1 signaling pathway | 24/421 | 2.2E-04 | 9.2E-03 | 24 |
| hsa04010 | MAPK signaling pathway | 29/421 | 6.3E-04 | 1.7E-02 | 29 |
| hsa05100 | Bacterial invasion of epithelial cells | 11/421 | 2.0E-03 | 3.2E-02 | 11 |
