## Supplemental Figure 1-17 for "High Resolution Single Cell Maps Reveals Distinct Cell Organization and Function Across Different Regions of the Human Intestine"

Supplemental Figures 1-17

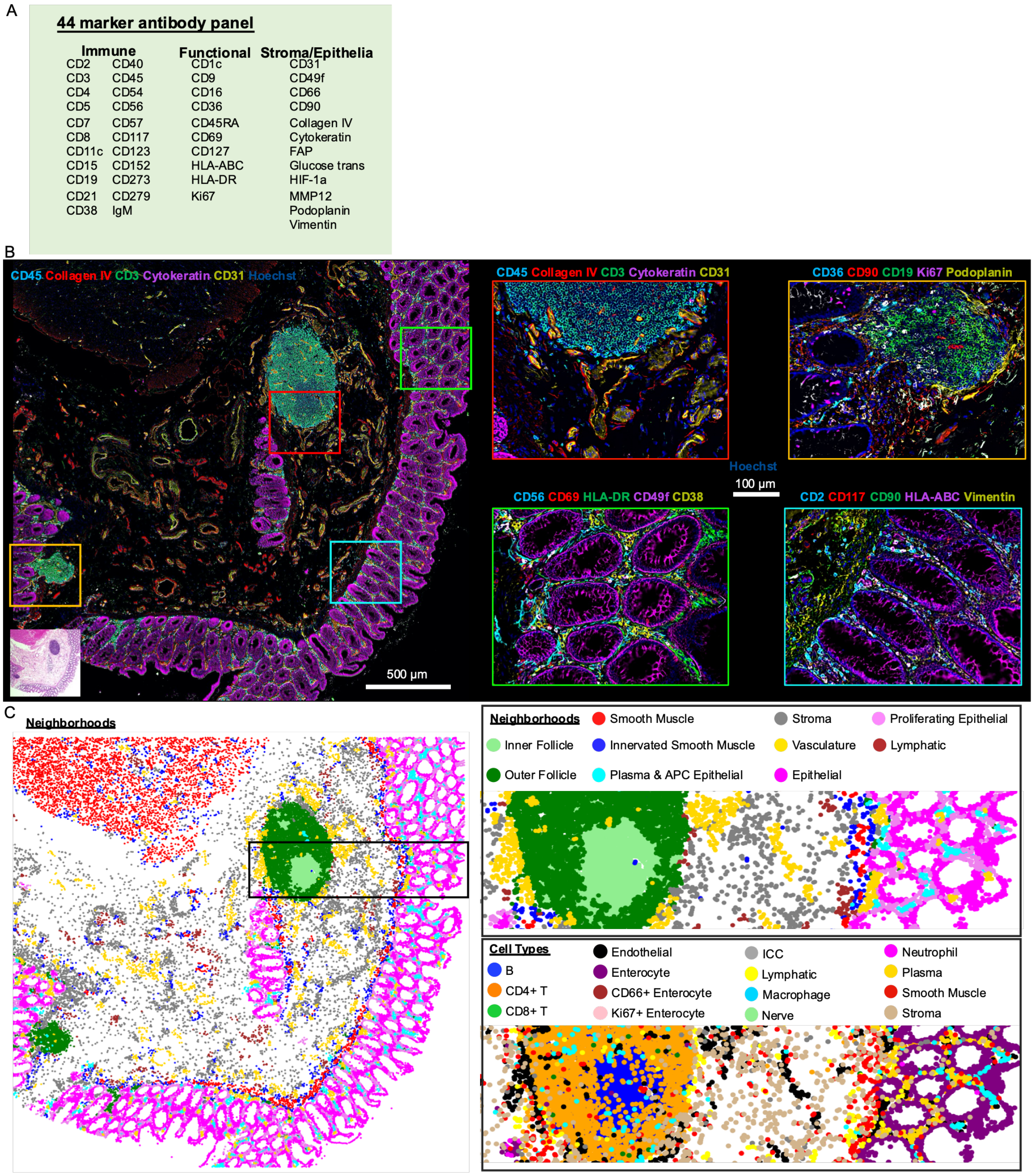


**Supplemental Figure 1:** CODEX multiplexed imaging of first donor B001 for small intestine and colon samples. A) 44 marker antibody base panel used for imaging. B) CODEX imaging of one of the 8 different sites taken from the colon for B001, with 5/44 markers shown. Four zoomed in regions with various 5 color combinations of markers from the larger image (denoted by colored outline) with also an H&E image shown for the tissue. C) Multicellular neighborhood map for the same region with both cell type and neighborhood maps zoomed in around immune follicle regions.

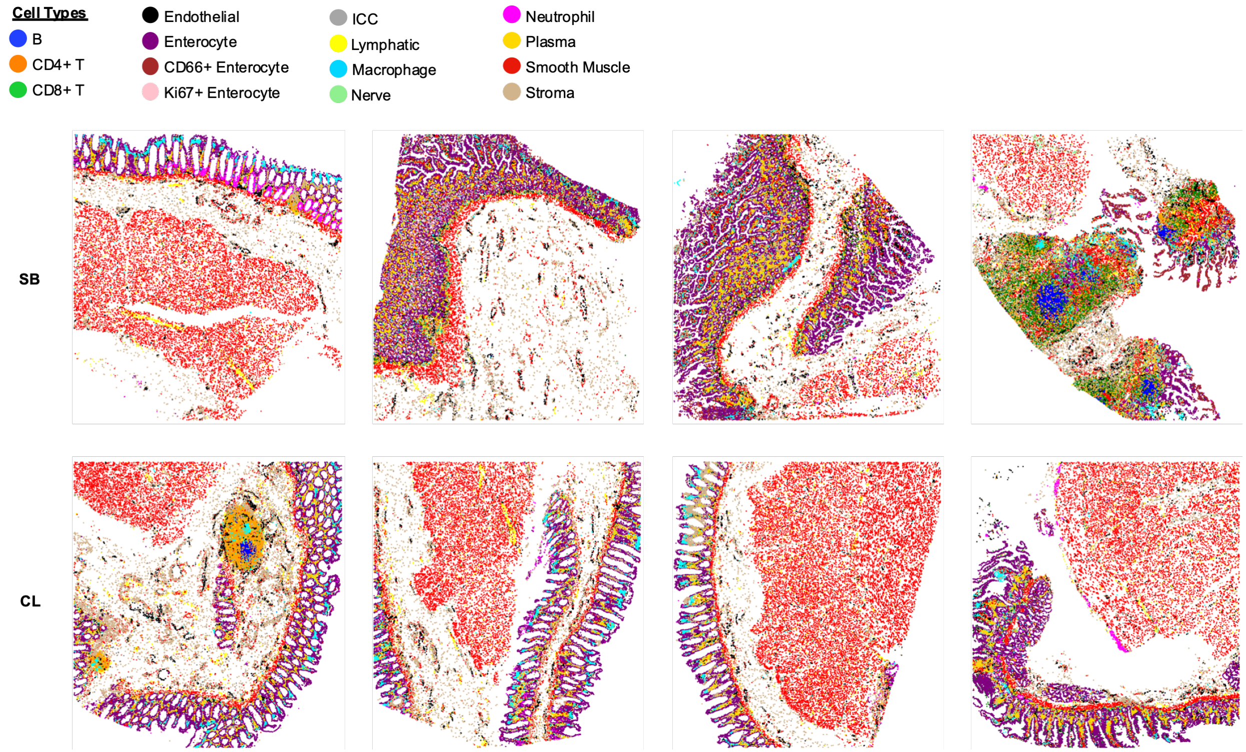


**Supplemental Figure 2:** Cell type maps for all 8 regions imaged for donor 1 (B001) of the small intestine and colon.

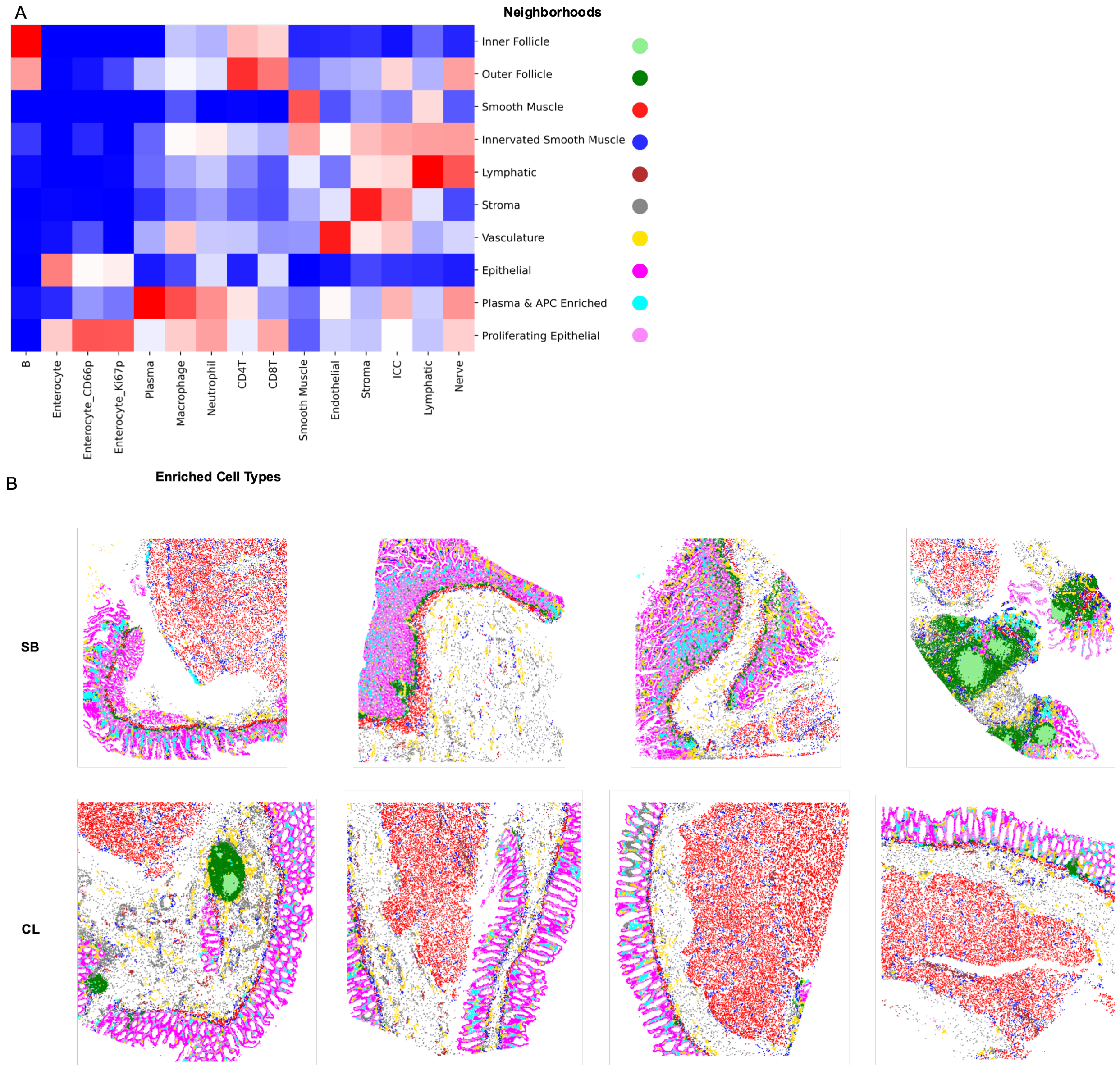


**Supplemental Figure 3:** Determination of multicellular neighborhoods in CODEX multiplexed imaging for donor 1 across all 8 regions. A) Cell type enrichment score within each neighborhood as represented in the heatmap. B) Cell type mapped back to each individual sample imaged.

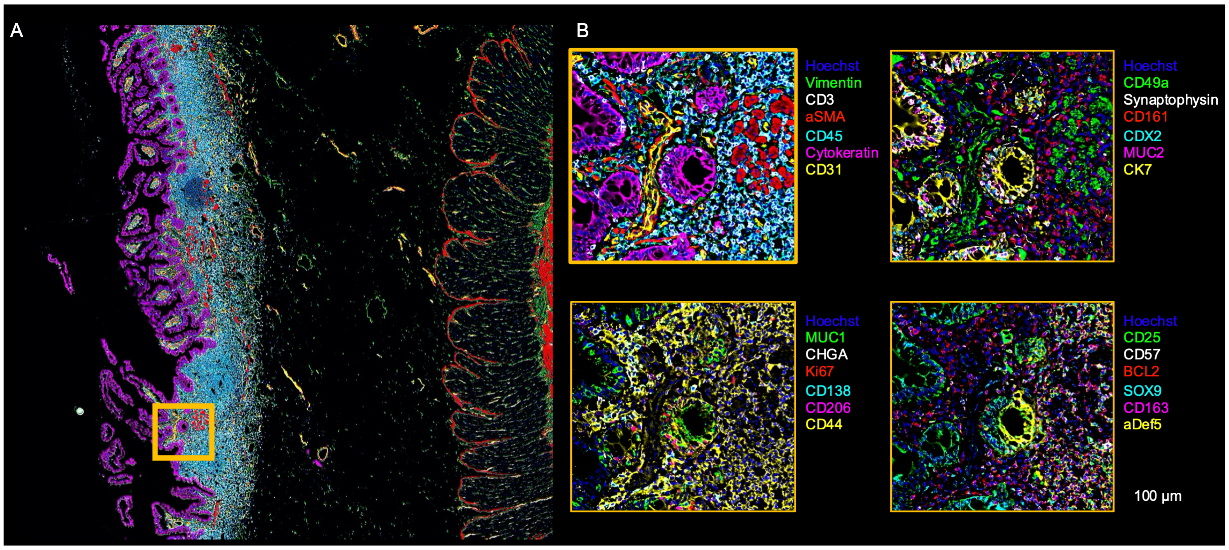


**Supplemental Figure 4:** CODEX multiplexed imaging with expanded 54 antibody CODEX panel. A) One region of B004 shown with 6 markers shown (Vimentin, CD3, aSMA, CD45, Cytokeratin, and CD31) B) Zoomed in regions with 6 markers each shown for the region highlighted (yellow box) in A with additional markers added to the panel highlighted.

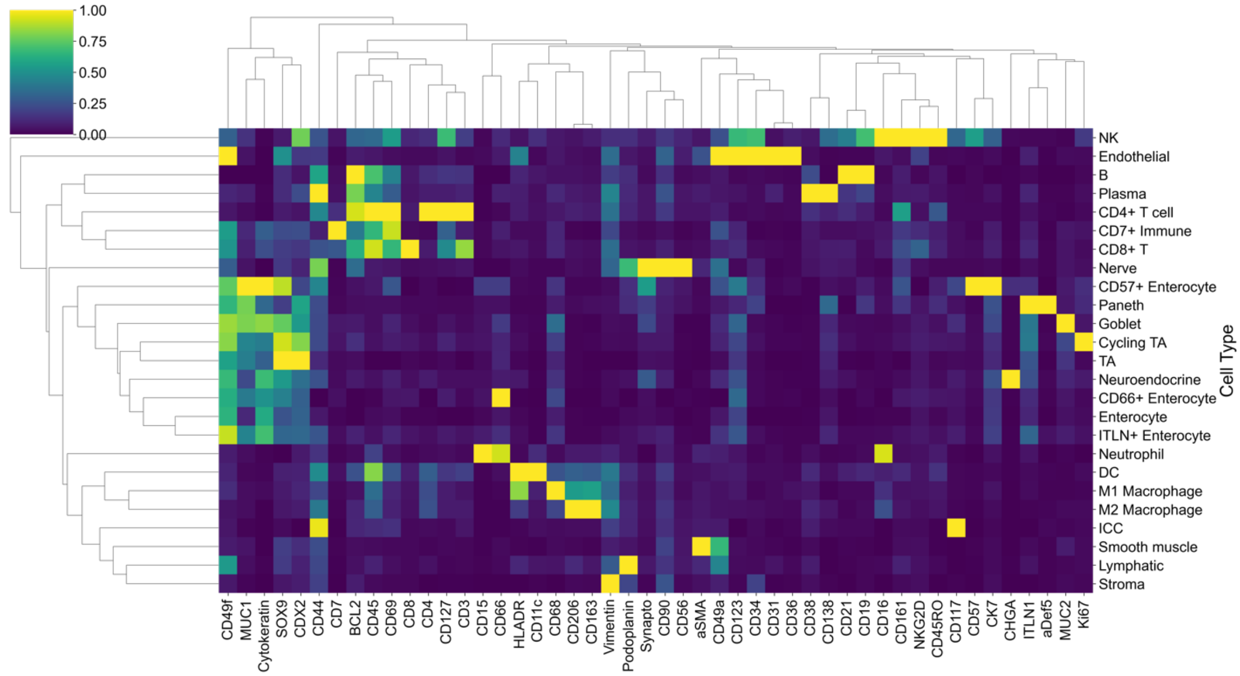


**Supplemental Figure 5:** Cell type by marker heatmap normalized both by column and row to show markers which define cell types from CODEX multiplexed imaging for samples B004,5,6.

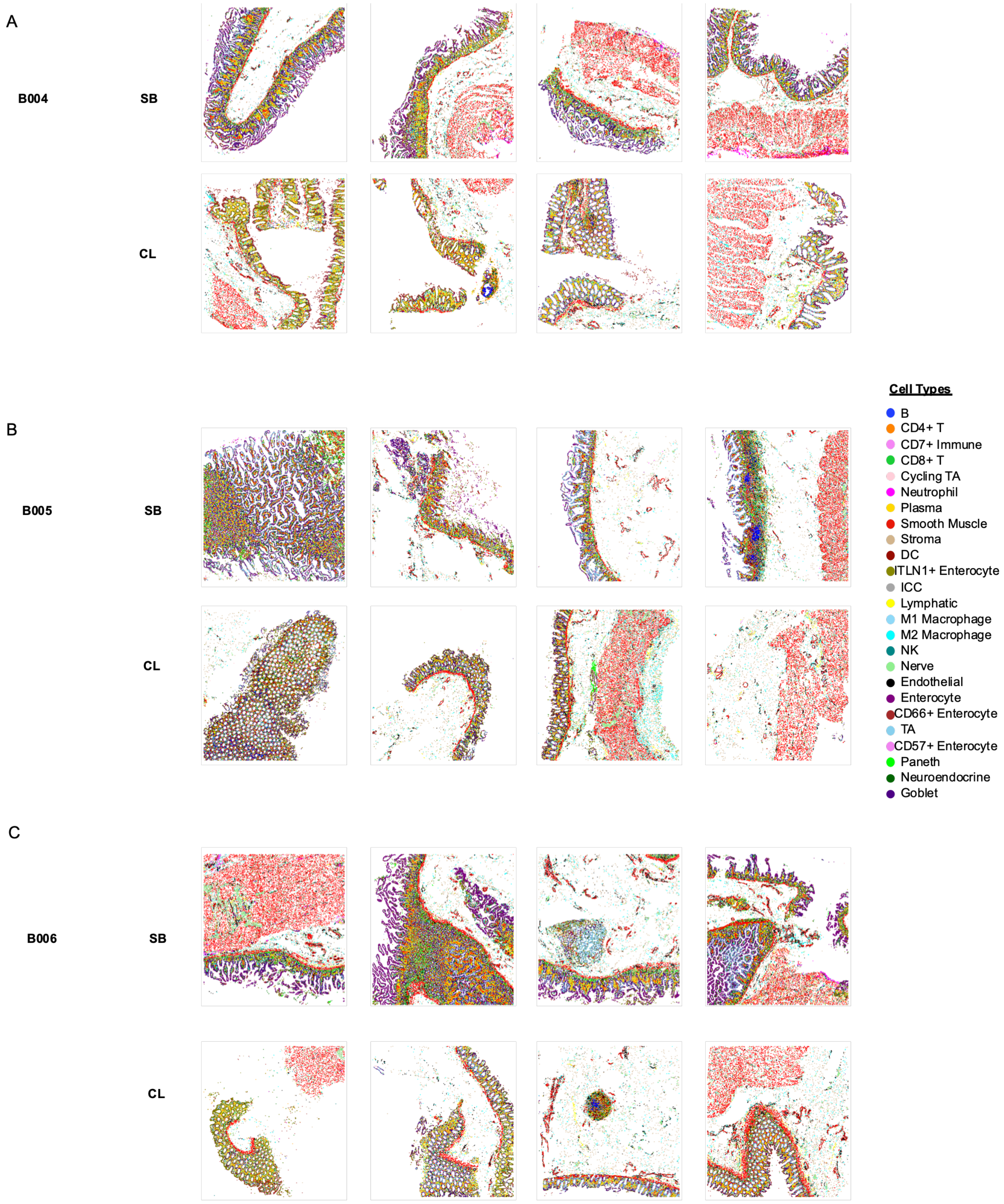


**Supplemental Figure 6:** Cell type maps for all 8 regions imaged for donors B004, 5, 6 of the small intestine and colon.

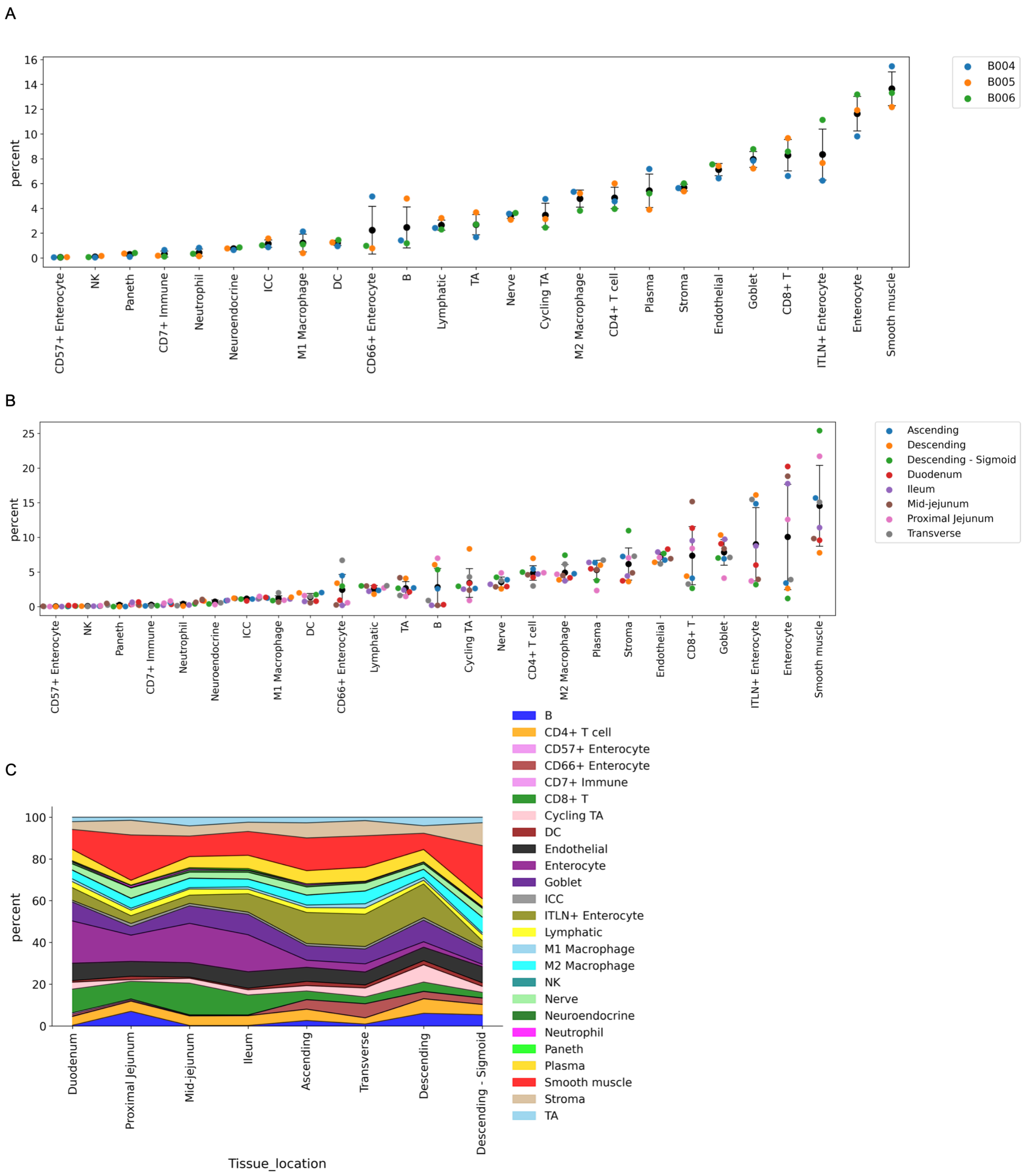


**Supplemental Figure 7:** Quantification of cell type percentages for B004,5,6. A) Cell type percentage separated by donor. B) Cell type percentage grouped by location in the intestine. C) Cell type percent summation graph by location in the intestine with all cell types shown.

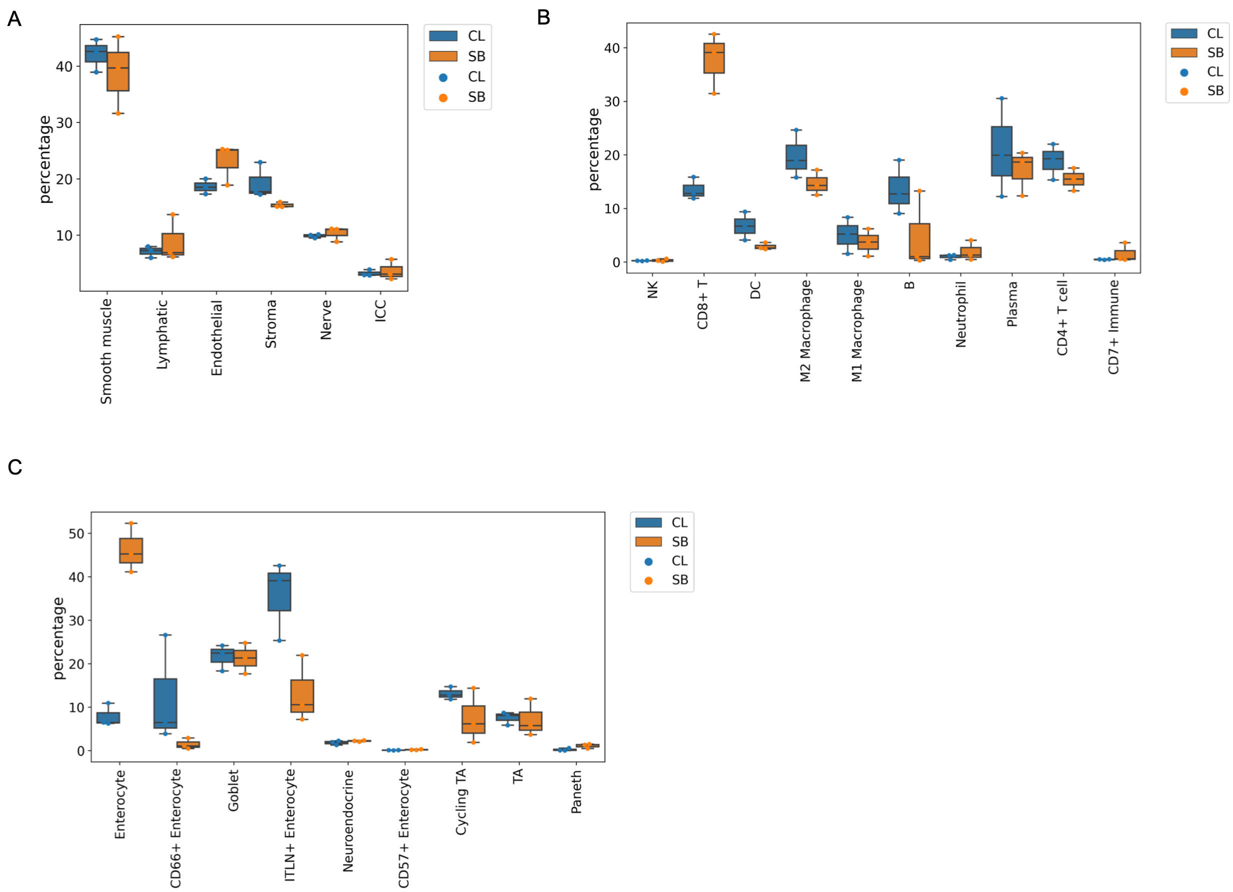


**Supplemental Figure 8:** Cell type percentages as determined by CODEX multiplexed imaging as separated within three categories: A) Stromal, B) Immune, and C) Epithelial types and compared for the colon (CL) and small bowel (SB).

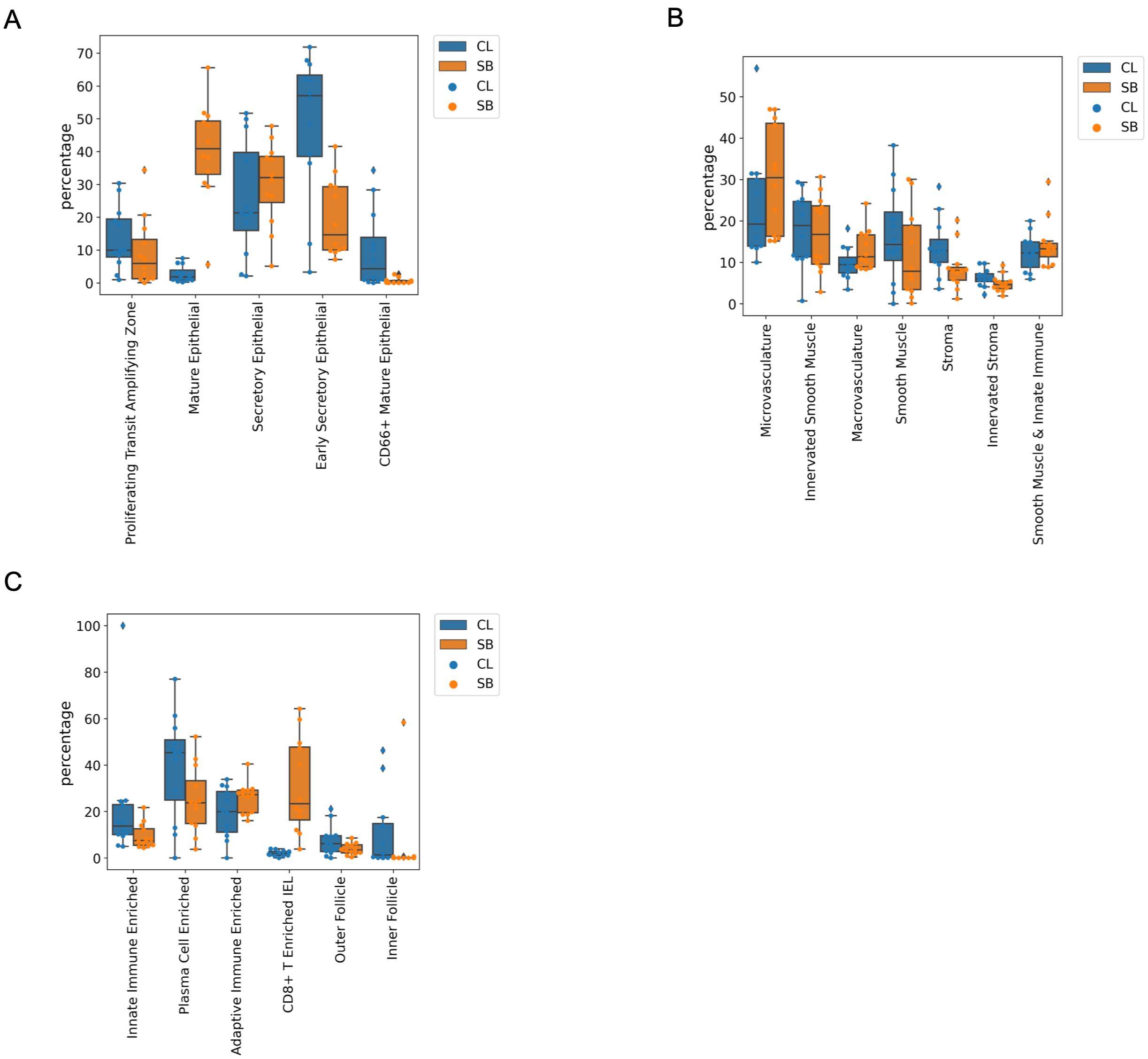


**Supplemental Figure 9:** Neighborhood type percentages as determined by CODEX multiplexed imaging as separated within three categories: A) Stromal, B) Immune, and C) Epithelial types and compared for the colon (CL) and small bowel (SB).

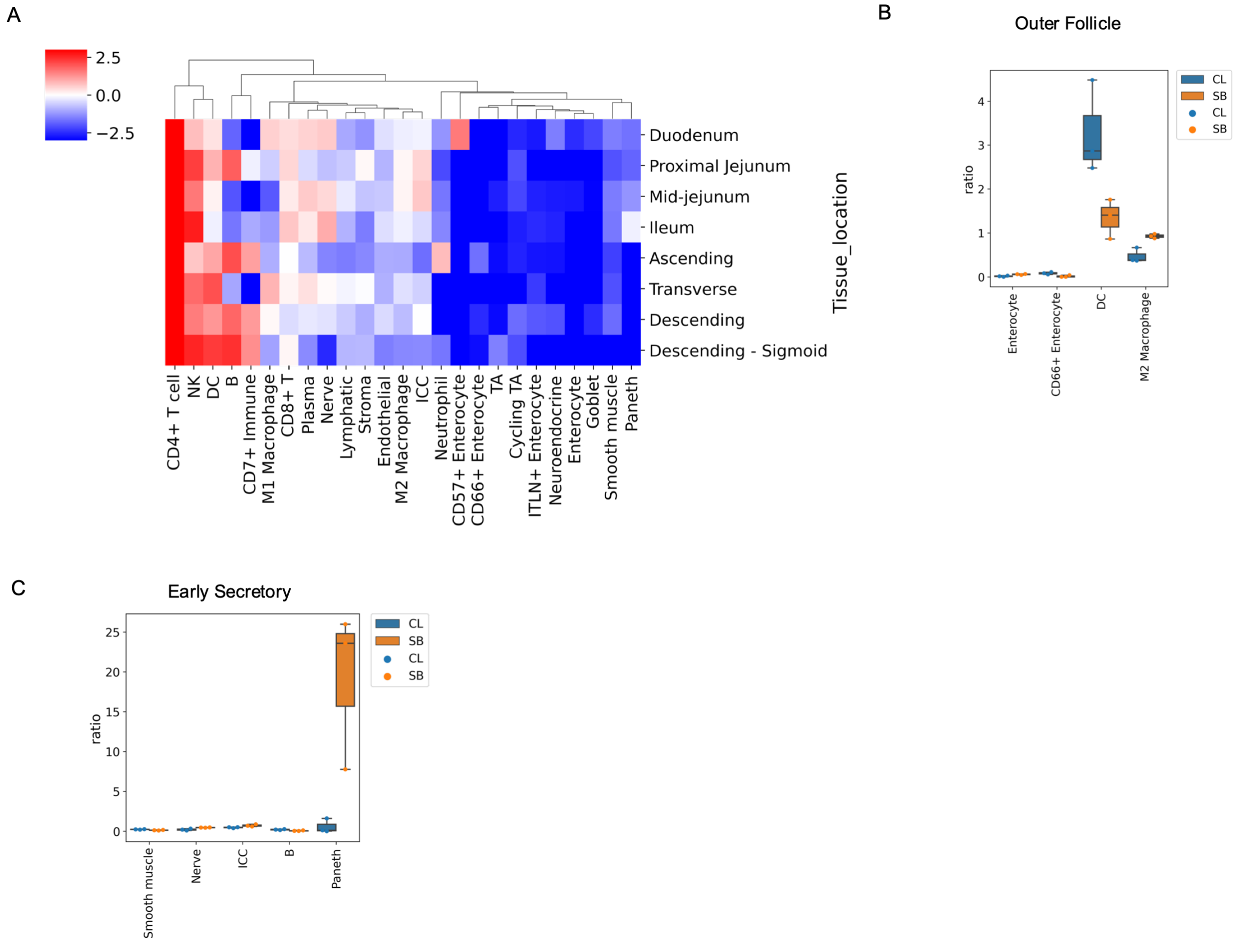


**Supplemental Figure 10:** Change in composition of cellular neighborhood across the intestine. A) Neighborhood *Outer Follicle* cell type enrichment for all cell types across the 8 regions measured from the intestine. B-C) Quantification of major cell types changed with quantified replicates shown for B) *Outer Follicle* neighborhood and C) *Early Secretory* neighborhood.

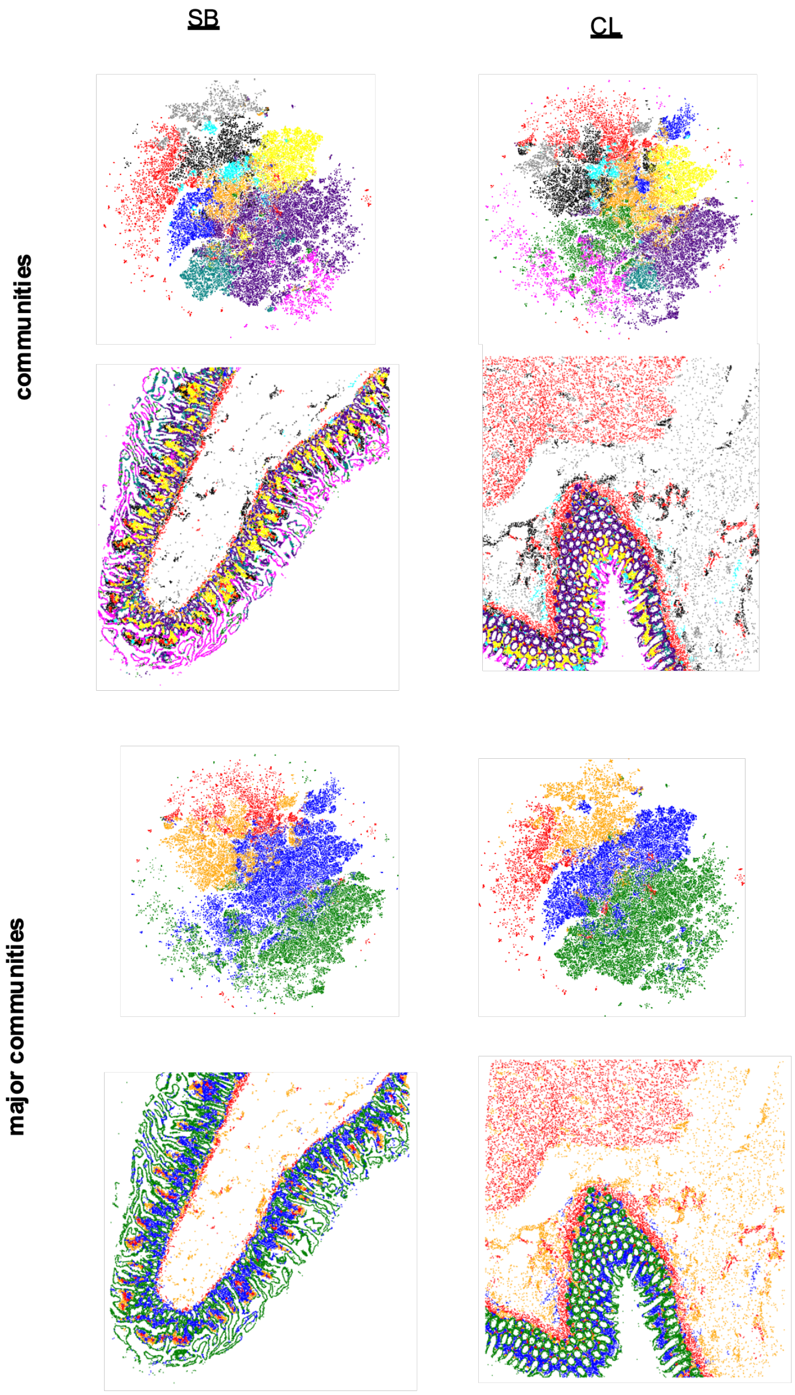


**Supplemental Figure 11:** Community and major community type maps for small bowel (SB) and colon (CL) with associated tsne maps generated from vectors of neighborhood composition.

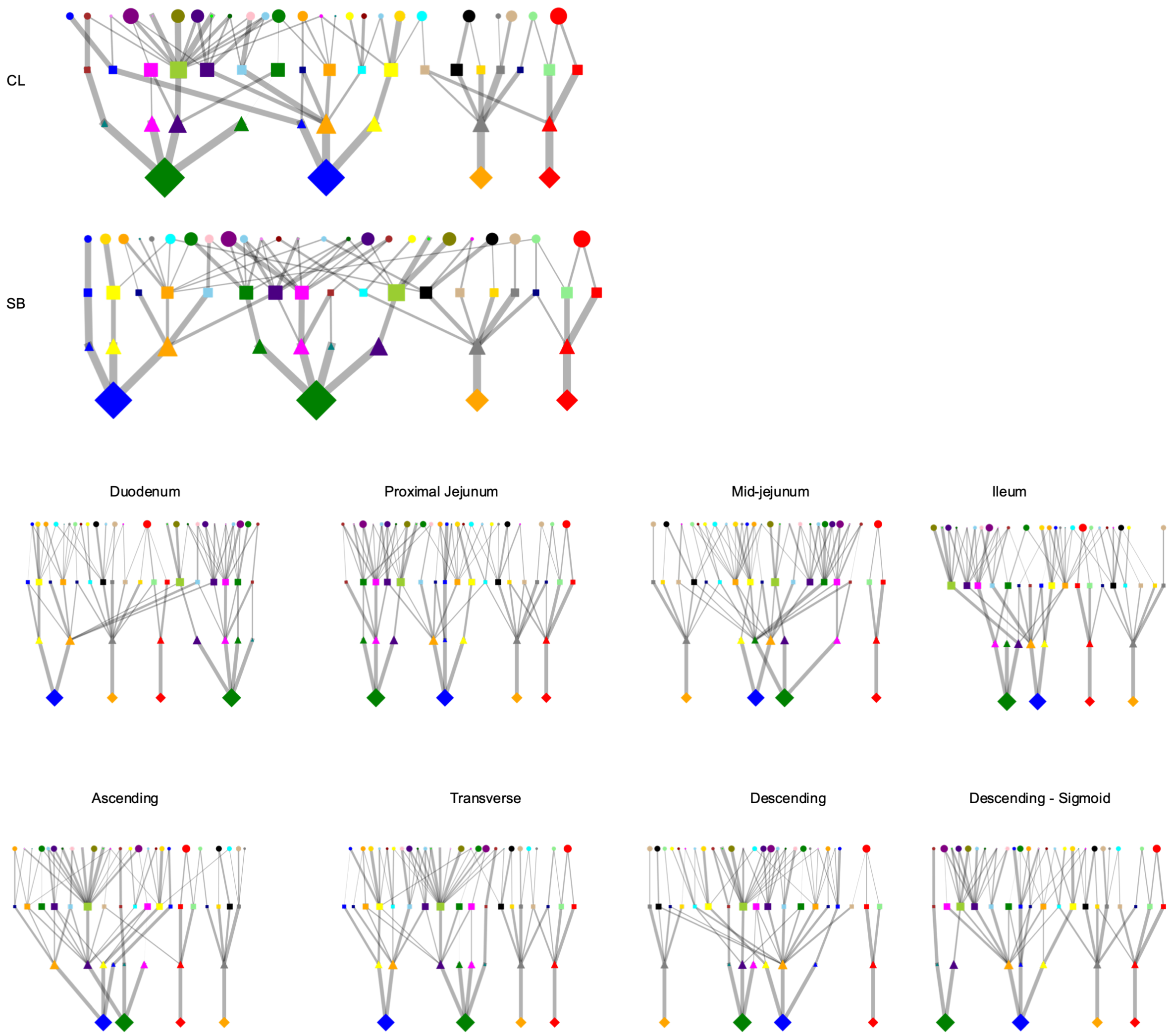


**Supplemental Figure 12:** Graph of multi-level structure of the tissue as broken down by the different structures. Shapes correspond to structural level, colors represent individual categories, size of shapes represents the percent contribution to tissue, and the size of connected lines represents the overall contribution to the next level of structure as moving down the graph in increasing tissue structural hierarchy. These are separated either by small bowel (SB) or colon (CL) and also by the eight different sites taken across the intestine.

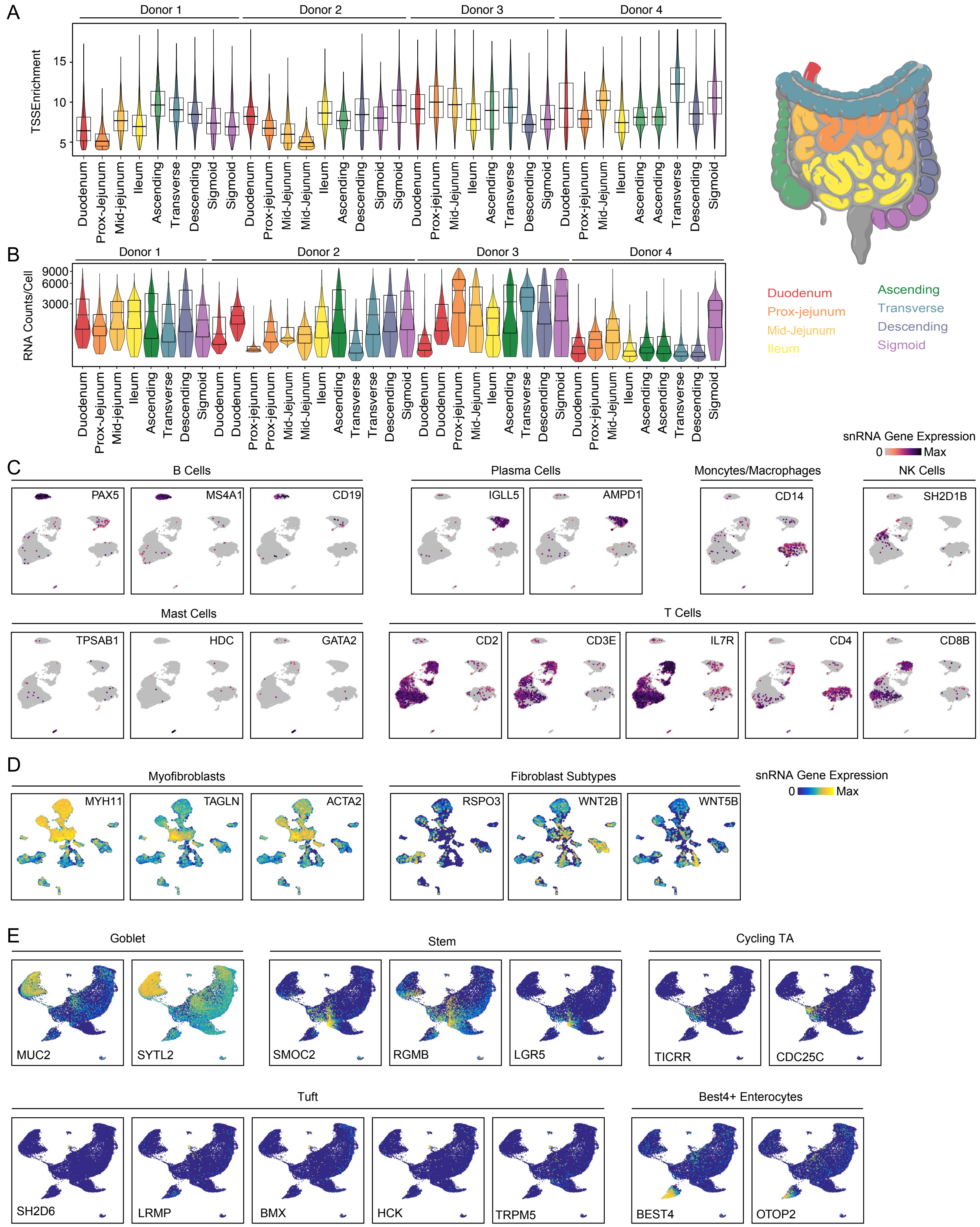


**Supplemental Figure 13:** Quality control and annotation of single-nuclei data.(A, B) Violin plots of TSS enrichment (A) and RNA counts/cell (B) for different samples included in the study. Samples are colored by the location from which they were obtained. (C) UMAP projection of immune snRNA cells colored by expression of marker genes. (D) UMAP projection of stromal snRNA cells colored by expression of marker genes. (E) UMAP projection of colon epithelial snRNA cells colored by expression of marker genes.

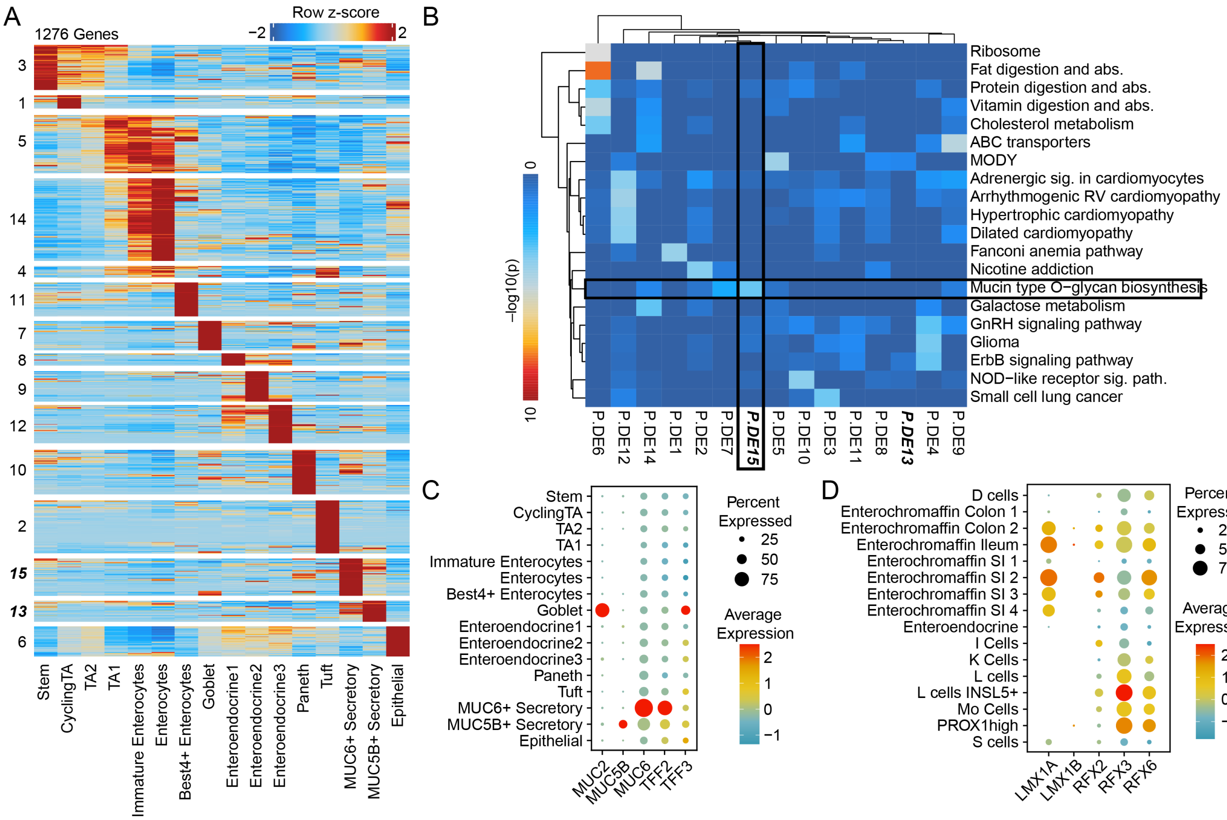


**Supplemental Figure 14:** Characterization of mucin secreting cell-types in the duodenum.(A) Heatmap representation of significantly differential genes between all cell types in the duodenum. Significantly differential genes were computed pairwise between all cell types in the duodenum. Genes that were differential between any two pair of cells were included in the set of differential gene. The resulting genes were kmeans clustered into 15 groups and the row z-score of expression of each gene is plotted in the heatmap.(B) Enrichment of kegg pathways in clusters of genes identified in A. The heatmap is colored by the –log10 of uncorrected p-values. (C) Dotplot representation of the expression of MUC2, MUC5B, MUC6, TFF2, and TFF3 in different cell types in the duodenum. (D) Dotplot representation of the expression of enteroendocrine/enterochromaffin TF regulators in enteroendocrine cell types.

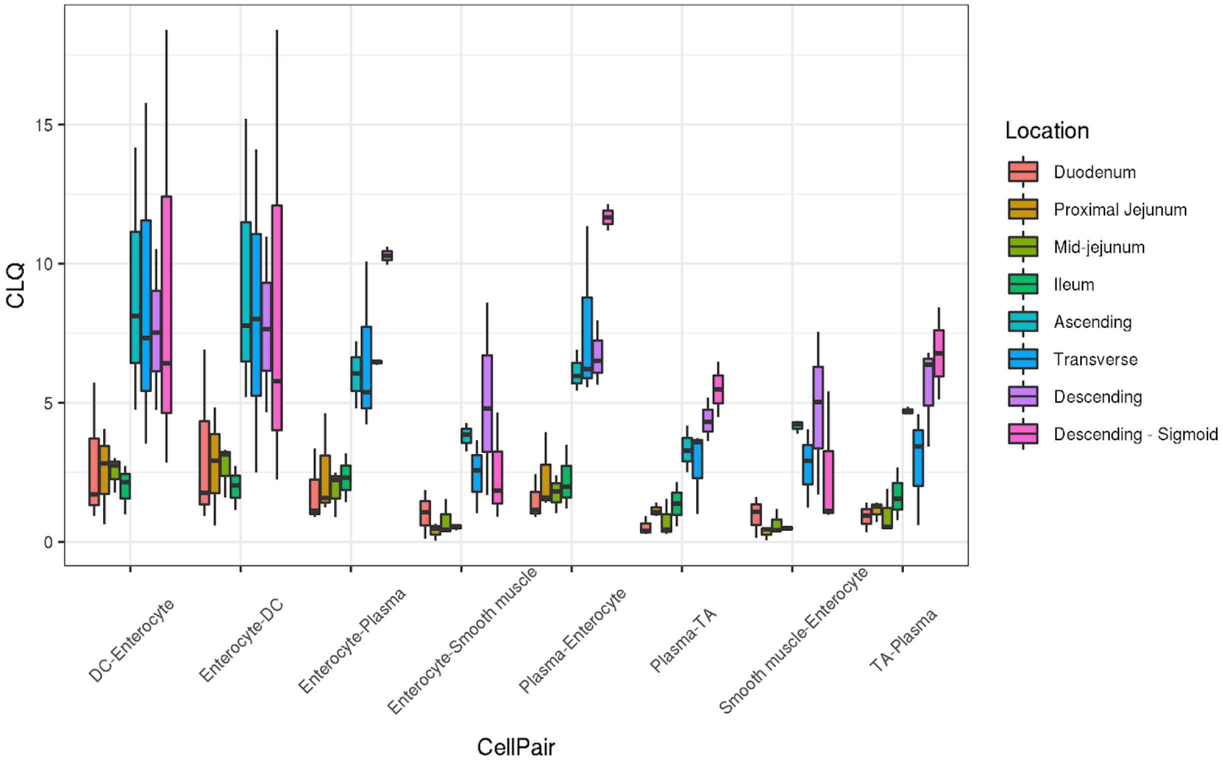


**Supplemental Figure 15:** Differences in pairwise cell-type colocalization patterns across tissue locations.

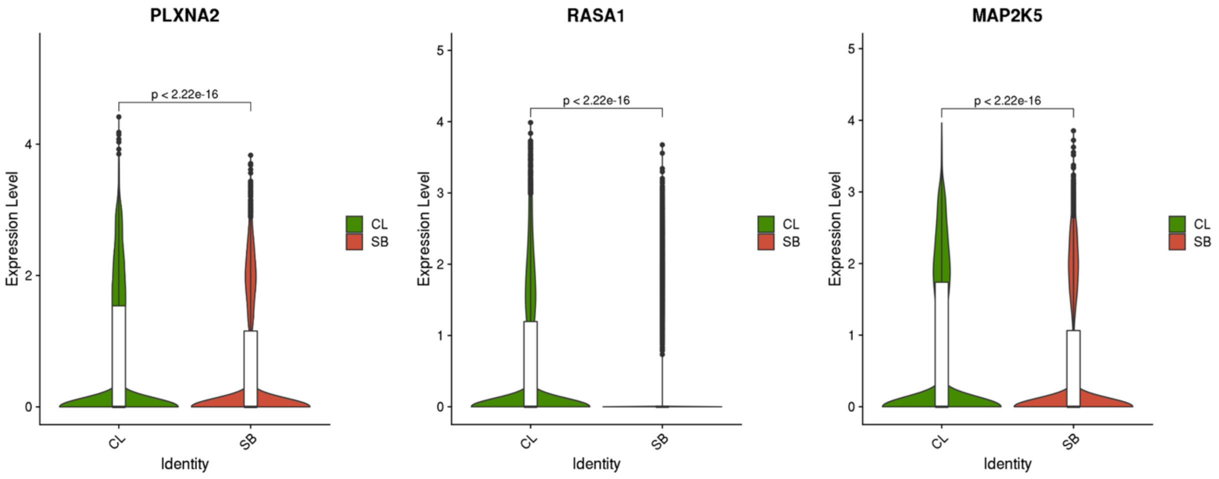


**Supplemental Figure 16:** PLXNA2, RASA1, and MAP2K5 expression in TA2 cells in large colon (CL) and small bowel (SB).

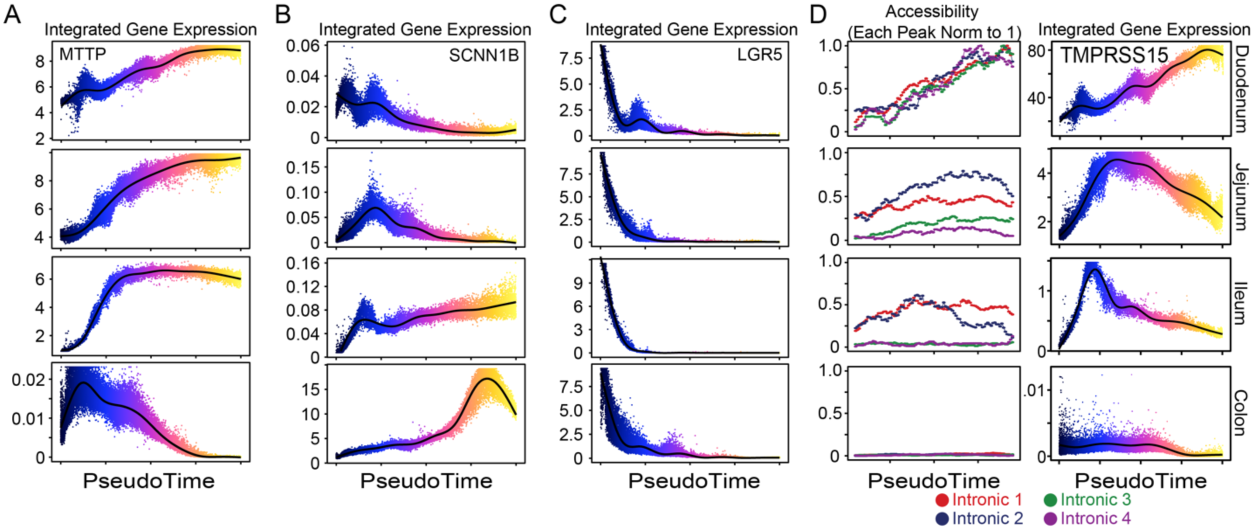


**Supplemental Figure 17:** Single gene changes along the absorptive differentiation trajectory. (A, B) Integrated gene expression of MTTP (A) and SCNN1B (B) and LGR5 (C) along the differentiation trajectory. (C) Accessibility at peaks correlated with the expression of TMPRSS15 along the differentiation trajectory in each region is plotted on the left. Each peak is normalized to the maximum accessibility along any of the trajectories. Integrated gene expression of TMPRSS15 along the differentiation trajectory in each region is plotted on the right.
